# Mitotic single-stranded DNA suppression by DDIAS

**DOI:** 10.1101/2025.09.09.675108

**Authors:** Yibo Xue, Faisal Bin Rashed, Daniel Y. L. Mao, Kento T. Abe, Zhen-Yuan Lin, Dheva Setiaputra, Lisa Hoeg, Clara Bonnet, Frank Sicheri, Anne-Claude Gingras, Daniel Durocher

**Affiliations:** Lunenfeld-Tanenbaum Research Institute, Mount Sinai Hospital, 600 University Avenue, Toronto, ON M5G 1X5, Canada; Department of Molecular Genetics, University of Toronto, 1 King’s College Circle, Toronto, ON M5S 1A8, Canada; Department of Biochemistry, University of Toronto, 1 King’s College Circle, Toronto, ON M5S 1A8, Canada

## Abstract

Incomplete DNA replication and chromosome breakage during mitosis pose major threats to chromosome segregation. The CIP2A-TOPBP1 complex acts to mitigate this peril, but its exact role is not yet understood. Here, we report that DDIAS acts as a DNA-binding effector of the CIP2A-TOPBP1 complex. DDIAS directly interacts with TOPBP1 and the disruption of this interaction or inactivation of its single-stranded DNA (ssDNA)-binding ability impairs genome integrity and causes synthetic lethality with BRCA1 and BRCA2 deficiency. DDIAS does not promote the putative end-tethering function of CIP2A-TOPBP1 but rather acts to suppress ssDNA during mitosis. This role is emphasized by a pronounced genetic interaction between the genes coding for DDIAS and DNA polymerase (. We conclude that DDIAS defines a mitotic DNA damage response that mitigates the threat of mitotic ssDNA arising from DNA replication stress, which we infer is a liability to accurate chromosome segregation. (144 words).

## Introduction

The accurate segregation of chromosomes is fundamental to cellular homeostasis, tumor suppression and heredity. Chromosome segregation occurs during mitosis, a short period of the cell cycle in which condensed chromosomes attach to bipolar spindles^1^ and then begin a poleward migration that requires the removal of DNA and protein linkages between sister chromatids^2^. The presence of DNA double-stand breaks (DSBs), under-replicated DNA^3^ and unresolved recombination intermediates^4,5^ at the time of cell division all pose major challenges to the accuracy of chromosome segregation. In the first case, chromosome breaks generate acentric chromosome fragments that have a high potential for mis-segregation due to their lack of attachment to the mitotic spindle. In the other two cases, under-replicated DNA or lingering recombination intermediates link sister chromatids together which can impair chromosome segregation or even result in mitotic failure. To counteract these challenges, cells have mitosis-specific pathways that mitigate the impact of DNA lesions and that promote the resolution of chromatid linkages^6^.

The recently described CIP2A-TOPBP1 complex is one such mitotic DNA damage response system^7–10^. CIP2A resides primarily in the cytoplasm during interphase. Following nuclear envelope breakdown, it forms a complex with the adaptor protein TOPBP1, and together they localize to mitotic DNA lesions, which include DSBs and under-replicated DNA. At mitotic DSBs, CIP2A-TOPBP1 is recruited in a chromatin- and MDC1-dependent manner where it may act as a chromosome end-tether that promotes biased (i.e. faithful) acentric chromosome fragment segregation^7,8^. One manifestation of this function can be found in the observation that CIP2A promotes the clustering and *en masse* segregation of fragmented micronuclear chromosomes during mitosis^11–13^. While this end-tethering role improves the odds of accurate acentric chromosome fragment segregation, it also promotes the genesis of chromothriptic chromosome rearrangements^13^.

CIP2A-TOPBP1 also localizes to DNA lesions that are formed as a consequence of incomplete DNA replication in a manner that is at least partly MDC1-independent^7^. Under-replication is a major challenge to cell division and a dedicated mitotic system is in place to process unreplicated chromatids and ensure timely chromatid disjunction. This system involves the CDK1-dependent activation of the replisome-associated TRAIP E3 ubiquitin ligase^14,15^ in complex with TTF2^16,17^, which ubiquitylates replisome components to promote disassembly and nucleolytic processing of the joined sister chromatids by one or more nucleases of currently unknown identity^14,15^. It is not yet clear what the function of CIP2A-TOPBP1 is at those lesions and whether it is recruited prior to or subsequent to replisome disassembly, or before or after nucleolytic processing. This role of CIP2A-TOPBP1 at under-replicated loci is likely the basis for the highly penetrant synthetic lethality between *CIP2A* and *BRCA1/2*^7,18^ (or *PALB2*^19^), as mitotic DNA lesions due to under-replication accumulate in BRCA-deficient cells^7,20^. Loss of CIP2A, or disabling the CIP2A-TOPBP1 interaction, in homologous recombination (HR)-deficient cells causes high levels of micronucleation characterized by acentric chromosome mis-segregation^7^. Whether this phenotype is due to defective end-tethering or whether it reflects defective repair or resolution of DNA lesions is currently unknown, although recent work^21^ suggests that CIP2A may also promote additional mitotic DNA damage response processes such as MiDAS or Polθ-mediated end-joining (TMEJ), a DSB repair pathway that is activated and prominent in mitotic cells^22–24^.

In this study, we aimed to further define the role of the CIP2A-TOPBP1 complex in the mitotic DNA damage response by searching for additional components of the pathway. Since TOPBP1 acts as a protein scaffold in multiple genome maintenance processes^25^, we considered the possibility that TOPBP1 may bridge CIP2A to other proteins through its ability to bind to phosphoproteins with its tandem BRCT domains^26^. This led us to discover that the function of CIP2A-TOPBP1 in promoting the viability of homologous recombination (HR)-deficient cells involves an interaction with the OB-fold containing protein DDIAS. We present evidence that DDIAS uses its ssDNA-binding activity to limit ssDNA during mitosis, which we propose represents a source of mitotic chromosome breakage and a liability in HR-deficient cells.

## Results

### The TOPBP1 C-terminus is essential in BRCA2-deficient cells

To assess whether any domain of TOPBP1 (Fig 1a), other than its CIP2A-binding motif, promotes the CIP2A-dependent mitotic DNA damage response, we developed a conditional gene replacement system in which *TOPBP1^-/-^* cells generated by gene editing (Fig. S1a,b and Table S1) are kept alive through a doxycycline-repressible transgene expressing TOPBP1 fused to the degron mAID (producing mAID-TOPBP1; the cell line also expresses TIR1-F74G^27^; Fig. S1a,b). We generated an isogenic variant of this cell line in which *BRCA2* was inactivated by gene editing (*BRCA2-KO.1*; Fig S1c,d and Table S1). Depletion of mAID-TOPBP1 upon the combined addition of doxycycline and 5-Ph-IAA, a derivative of the auxin hormone indole-3-acetic acid (the “DI” condition), was lethal, which was expected since *TOPBP1* is an essential gene (Fig. 1b,c, EV condition). We transduced the two *mAID-TOPBP1* cell lines with lentivirus expressing mutants that expressed wild-type (WT) HA-tagged TOPBP1 or variants that disabled BRCT1 (K154A), BRCT1/2 (K154A/K155A/K250A; referred to as 3KA), BRCT3 (Δ354-444), the BRCT4/5 module (K704A), BRCT6 (Δ901-985) and the BRCT7/8 module (K1317A) (Fig 1a). We also expressed TOPBP1-F1071A, which abrogates condensate formation by TOPBP1^28^ and the F837A-D838A-V839A mutant (referred to as 3A) that disrupts interaction with CIP2A^7^. All the TOPBP1 variants expressed at levels similar to those of TOPBP1 WT (Fig S1e). Next, we inactivated mAID-TOPBP1 and examined micronucleation (MN) as a surrogate for genomic instability in both the BRCA2-proficient and -*KO.1* cell lines (Fig S1f). In BRCA2-proficient cells, loss of TOPBP1 caused high levels of micronucleation that were completely suppressed by the lentiviral expression of TOPBP1 WT (Fig 1d and Fig S1g). Only two TOPBP1 mutants caused an increase in micronucleation in the presence of BRCA2: the 3KA and the Δ354-444 mutants (Fig 1d). The same two mutants were the only mutants that caused lethality in BRCA2-proficient cells after mAID-TOPBP1 inactivation (Fig 1b,c). Given that the BRCT0/1/2 module binds to Treslin (also known as TICRR)^29,30^, and BRCT3 binds to DONSON^31^, we surmise that the impaired DNA replication caused by the loss of these interactions is responsible for the micronucleation and cell lethality phenotypes.

**Figure 1.**
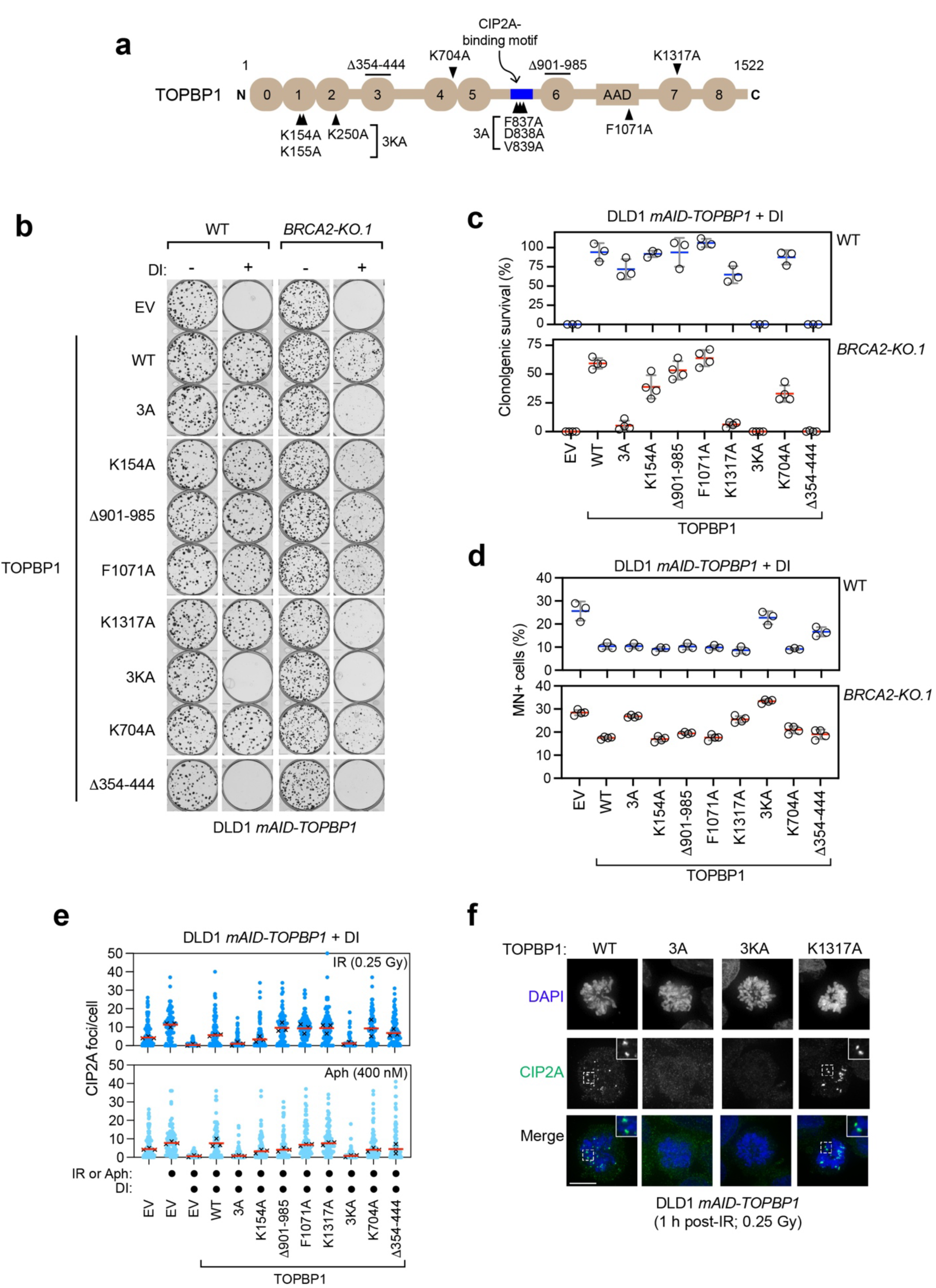
The TOPBP1 BRCT7/8 module is essential in BRCA2-deficient cells. (**a**) Schematic showing domain structure of the TOPBP1 protein. Numbered circles represent BRCT domains. AAD: ATR-activating domain. (**b, c**) Clonogenic survival of DLD1 *mAID-TOPBP1* WT and *BRCA2*^-/-^ cells expressing indicated the *TOPBP1* variants or EV (empty vector), followed by depletion of mAID-TOPBP1 by DI. DI: doxycycline (1 μg/mL) + 5-Ph-IAA (1 μM). Data was normalized to untreated condition. Bars indicate the mean ± SD. n=3 independent experiments for WT cells and n= 4 for *BRCA2*^-/-^ cells. (**d**) Quantification of micronuclei in cells (as in **b, c**) after treatment with DI for 72h. (**e**) Immunofluorescence analysis of CIP2A foci in mitotic DLD1 *mAID-TOPBP1* cells expressing indicated *TOPBP1* variants or EV, followed by depletion of mAID-TOPBP1 by DI. Cells were fixed 1h after 0.25 Gy IR treatment or 16h after 400nM aphidicolin (Aph) treatment. Blue dots represent measurements from individual cells and x marks represent the medians from three independent experiments. n=3 independent experiments (**f**) Representative micrographs of the experiment shown in **e**. Scale bar, 10 μm.

In parallel, we profiled the same set of TOPBP1 variants in *mAID-TOPBP1 BRCA2-KO.1* cells, assessing both micronucleation and viability in clonogenic survival assays. We found that the TOPBP1-3A and -K1317A mutants displayed an increase in micronucleation that was specific to BRCA2-deficient cells (Fig 1d and Fig S1g), and this higher level of genomic stability was accompanied by *BRCA2* mutant-specific cell lethality (Fig 1b,c). Therefore, similar to the loss of the CIP2A-TOPBP1 interaction, which is disabled by the 3A mutation^7^, loss of phosphopeptide-binding imparted by the K1317A causes synthetic lethality with BRCA2 deficiency. These results suggest that TOPBP1 interacts with one or more proteins via its BRCT7/8 domain that may also be involved in the CIP2A pathway.

Since the TOPBP1-CIP2A interaction is critical for the recruitment of each protein into mitotic DNA damage foci, we monitored the impact of the TOPBP1 mutants described above on the localization of CIP2A to mitotic DNA lesions induced either by X-irradiation (IR; 0.25 Gy) or after low-dose aphidicolin treatment (Aph; 400 nM) for 16 h. As expected, and unlike TOPBP1 WT, the TOPBP1-3A mutant is defective in forming IR- or Aph-induced CIP2A foci, as previously reported^7^ (Fig 1e,f and Fig. S2a). Similarly, disruption of the N-terminal BRCT0/1/2 modules, in particular by the 3KA mutation, impairs CIP2A recruitment to mitotic DNA lesions (Fig 1e,f and Fig. S2a). This result is consistent with the observation that CIP2A-TOPBP1 recruitment to IR-induced mitotic DSB sites depends on MDC1^7,8^, whose phosphorylated form is bound by the BRCT0/1/2 module^9^. In contrast, the K1317A mutation, which is lethal in *BRCA2^-/-^* cells, does not impair CIP2A recruitment to DSB sites (Fig 1e,f and Fig. S2a). To confirm these findings, we introduced the K1317A mutation at the endogenous *TOPBP1* locus in DLD1 cells (Fig S2b) and found that neither CIP2A nor TOPBP1 had impaired localization to mitotic DNA damage in *TOPBP1-K1317A* cells (Fig S2c,d). These data further support the idea that the TOPBP1 BRCT7/8 module interacts with a phosphorylated protein that acts downstream of CIP2A-TOPBP1.

Given the involvement of CIP2A-TOPBP1 in the tethering of shattered micronuclear chromosomes^11–13^, we tested whether the TOPBP1-K1317A mutation impaired chromosome fragment tethering. We inserted a sequence encoding a dTAG conditional degron^32^ at the C-terminus of endogenous TOPBP1 (Fig S3a-c) in a DLD1 derivative engineered to selectively inactivate the *Chr Y* centromere (known as CEN-Select)^33,34^, which causes high-levels of Y chromosome-specific micronucleation and micronuclear chromosome shattering upon the addition of doxycycline and IAA^11,13^. We lentivirally expressed HA-TOPBP1 and its 3A and K1317A derivatives in this cell line and, as previously reported^11–13^, we found that depletion of TOPBP1 caused micronuclear chromosome dispersion in mitotic cells, a phenotype that could be completely reversed by expression of WT TOPBP1 (Fig S3d-g). Conversely, expression of the TOPBP1-3A mutation was unable to rescue micronuclear chromosome dispersion, consistent with the requirement for CIP2A in this process^11–13^ (Fig S3d-g). To our surprise, shattered micronuclear chromosomes were clustered efficiently in TOPBP1-K1317A-expressing cells (Fig S3d-g). These results indicate that the function of CIP2A-TOPBP1 in promoting viability of BRCA-deficient cells is distinct from its role in broken micronuclear chromosome tethering, and that these distinct roles can be genetically separated using the TOPBP1 K1317A mutation.

### Identification of DDIAS as a TOPBP1-interacting protein

Next, we initiated a search for proteins that interact with the CIP2A-TOPBP1 complex in a K1317-dependent manner. We developed an in-house variation of the split-TurboID system^35^ in which we fused an inactive N-terminal portion of miniTurbo (mT_N_) to the N-terminus of CIP2A and an inactive miniTurbo C-terminal fragment (mT_C_) to the C-terminus of TOPBP1 with the aim of re-forming an active miniTurbo following CIP2A-TOPBP1 complex formation in mitosis (Fig 2a). Addition of biotin to *BRCA2^-/-^* cells co-expressing mT_N_-Flag-CIP2A with HA-TOPBP1-mT_C_ (or HA-TOPBP1-K1317A-mT_C_) labeled mitotic foci that also contained CIP2A and TOPBP1 (Fig S3h). The biotin-containing foci were due to CIP2A-TOPBP1 complex formation since they were absent when a CIP2A-binding deficient^7^ TOPBP1-Δ756-891-mT_C_ fusion was used (Fig S3h). Cells expressing mT_N_-Flag-CIP2A in combination with either wild type HA-TOPBP1-mT_C_ or HA-TOPBP1-K1317A-mT_C_ were synchronized in mitosis and incubated with biotin for proximity labeling. The two cell lines were processed in parallel for biotinylated protein isolation (Fig S3i) that were then subjected to identification by mass spectrometry (Table S2). While most proteins identified, such as MDC1, were retrieved at similar levels in the WT and K1317A conditions (Fig 2b and Table S2), 7 proteins were identified in proximity of CIP2A-TOPBP1 complex in a K1317-dependent manner. Of those, BRIP1/FANCJ and PLK1, were two previously reported BRCT7/8-interacting proteins^36,37^ with BRCA1 being itself an interactor of BRIP1^38^. Of the remaining four, two were of particular interest: the above mentioned TTF2, a SNF2-family ATP-dependent translocase involved in TRAIP-dependent replisome unloading in mitosis mentioned above; and DDIAS (Fig 2c), a poorly characterized protein previously associated with apoptosis^39^ that co-evolved with the HR system^40^. DDIAS was especially interesting for two reasons: first, it had been identified in the same *BRCA1/BRCA2* synthetic lethality CRISPR screens that identified CIP2A^7^, hinting at a shared function; and second, it harbors an OB-fold domain (Fig 2c), which is often present in DNA binding-genome maintenance factors where it acts as a single-stranded (ss) DNA binding domain^41^. We validated that DDIAS, expressed as an HA-tagged fusion protein, can be co-immunoprecipitated by the isolated Flag-TOPBP1-BRCT7/8 module in a manner that depends on the TOPBP1 K1317 residue (Fig 2d). As controls we ascertained that both PLK1 and BRIP1 interact with TOPBP1-BRCT7/8 in a K1317-dependent manner, as expected (Fig 2d). DDIAS is therefore a TOPBP1-interacting protein.

**Figure 2.**
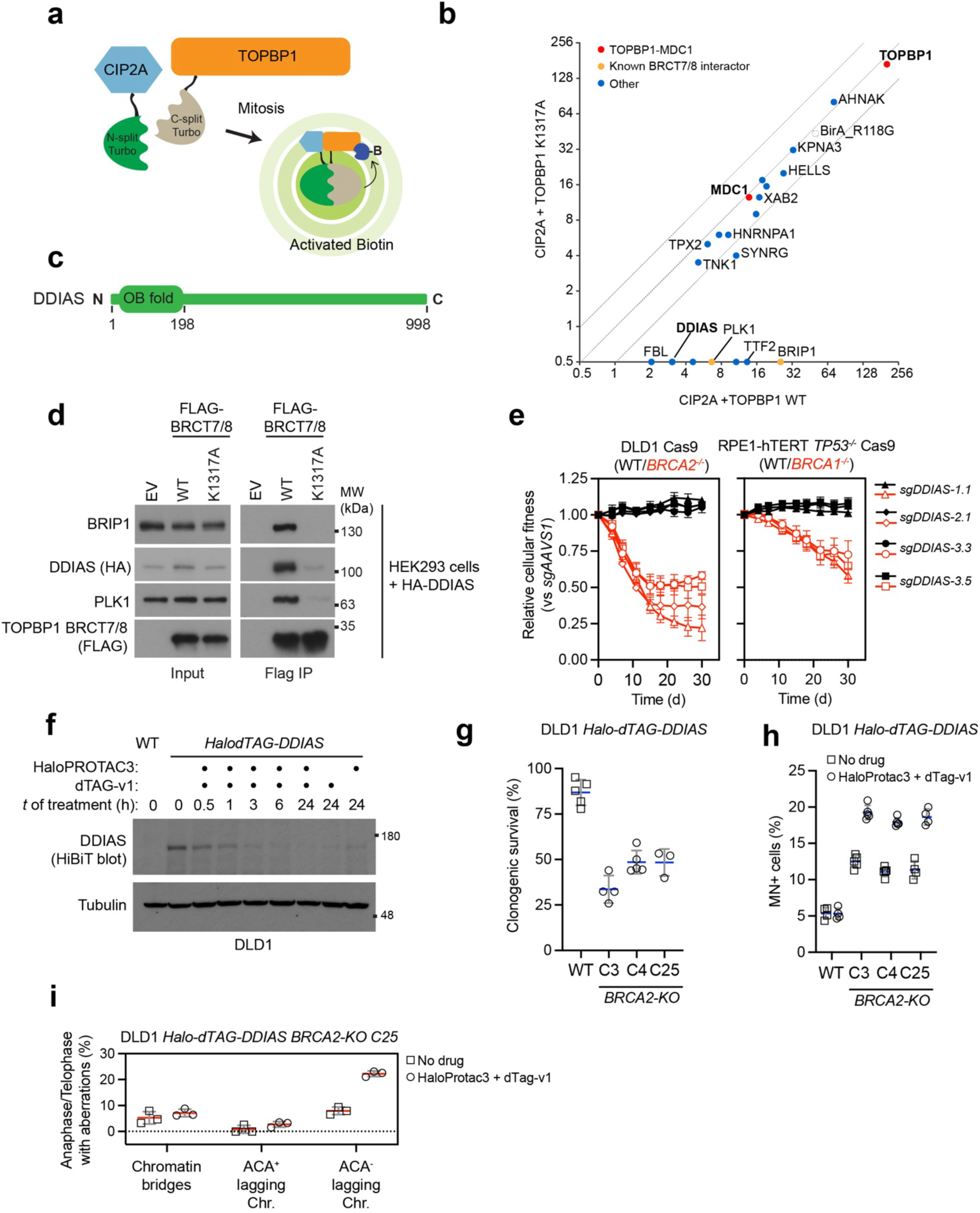
DDIAS interacts with the CIP2A-TOPBP1 complex. (**a**) Schematic of Split-TurboID with CIP2A and TOPBP1 tagged by the TurboID enzyme N-terminus and C-terminus, respectively. (**b**) Scatterplot showing average spectral counts from mass spectrometry analysis of CIP2A-TOPBP1 Split-TurboID. n=2 independent experiments. See Table S2 for raw data. (**c**) Schematic showing the domain structure of the DDIAS protein. (**d**) Immunoprecipitation of WT or K1317A-mutated Flag-BRCT7/8 in HEK293 cells transfected with HA-DDIAS. n=2 independent experiments. (**e**) Competitive growth assays in DLD1 Cas9 and RPE1-hTERT *TP53^-/-^* Cas9 cells with the indicated genetic background and the expression of the indicated sgRNAs. Data are presented as mean ± SD, normalized to day 0. n = 3 independent experiments. (**f)** HiBiT-Blot showing DDIAS degradation dynamics in DLD1 *HalodTAG-DDIAS* cells by HaloPROTAC3 (1 μM) and/or dTAG-v1 (1 μM). Tubulin, loading control. (**g, h)** Clonogenic survival (**g)** and micronuclei (**h)** in DLD1 *HalodTAG-DDIAS* cells with the indicated genetic background after DDIAS degradation. Cells were treated with HaloPROTAC3 (1 μM) + dTAG-v1 (1 μM) for 12 days in (**g)** and 3 days in (**h**). Clonogenic survival data was normalized to untreated condition. Bars indicate the mean ± SD, with n ≥3 independent experiments. (**i**) Quantification of chromosome segregation aberrations in anaphase or telophase of DLD1 *HalodTAG-DDIAS BRCA2*^-/-^ cells with or without DDIAS depletion. Lagging chromosomes were scored as containing (ACA^+^) or lacking (ACA^−^) a centromere detected with an anti-centromere antibody (ACA).

### DDIAS loss impairs viability of HR-deficient cells

To confirm that *DDIAS* and *BRCA1/2* display a synthetic-lethal relationship, we conducted cell competition assays that assessed the fitness of *BRCA1^-/-^* and *BRCA2^-/-^* cells following depletion of DDIAS with 3 (for *BRCA1^-/-^*) or 4 (for *BRCA2^-/-^*) independent single-guide (sg) RNAs. In line with the screen results, *DDIAS* inactivation impaired cell fitness selectively in the BRCA1/2-deficient cells (Fig 2e and Table S3). Targeting *DDIAS* with the same sgRNAs also reduced clonogenic survival in *BRCA1^-/-^*and *BRCA2^-/-^* cells, with the extent of the lethality approximately half of that caused by CIP2A loss^7^ (Fig S4a,b). As the project advanced, we generated a cell line enabling conditional inactivation of DDIAS with HaloPROTAC3 and dTAG-v1 (*HiBiT-HaloTag-dTAG-3xFLAG-DDIAS*, referred hereafter as *HalodTAG-DDIAS*) in DLD1 WT (Fig 2f and Fig S4c,d) and those in which we inactivated *BRCA2*, producing three *BRCA2-KO* clones (C3, C4 and C25; Fig S4e,f and Table S1). Sustained DDIAS degradation led to reduced clonogenic survival in the *BRCA2-KO* clones, accompanied by increased levels of micronucleation (Fig 2g,h and Fig S4g) and acentric lagging chromosomes (Fig 2i and Fig S4h). These results indicate that DDIAS promotes the viability and genome integrity of HR-deficient cells.

### DDIAS localizes to mitotic DNA damage sites downstream of CIP2A-TOPBP1

Next, we assessed whether DDIAS localizes at sites of mitotic DNA damage. We began these studies by exogenously expressing GFP-tagged DDIAS in DLD1 cells and their *BRCA2^-/-^*counterparts. We assessed the formation of DDIAS (i.e. GFP), TOPBP1 or γ-H2AX foci in mitotic cells that were previously treated with either aphidicolin (400 nM) or IR (0.25 Gy). GFP-DDIAS localization to spontaneous mitotic DNA lesions in *BRCA2^-/-^* cells was also monitored. In all cases, we observed that DDIAS co-localizes nearly perfectly with TOPBP1 at sites of mitotic DNA lesions marked by γ-H2AX (Fig 3a,b). This localization was specific to mitosis since we did not detect GFP-DDIAS foci in interphase (Fig 3b and Fig S5a). The GFP-DDIAS localization to IR-induced mitotic foci was completely dependent on CIP2A and TOPBP1 (Fig S5b,c) and was also dependent on the TOPBP1 K1317 residue (Fig 3c and Fig S5d).

**Figure 3.**
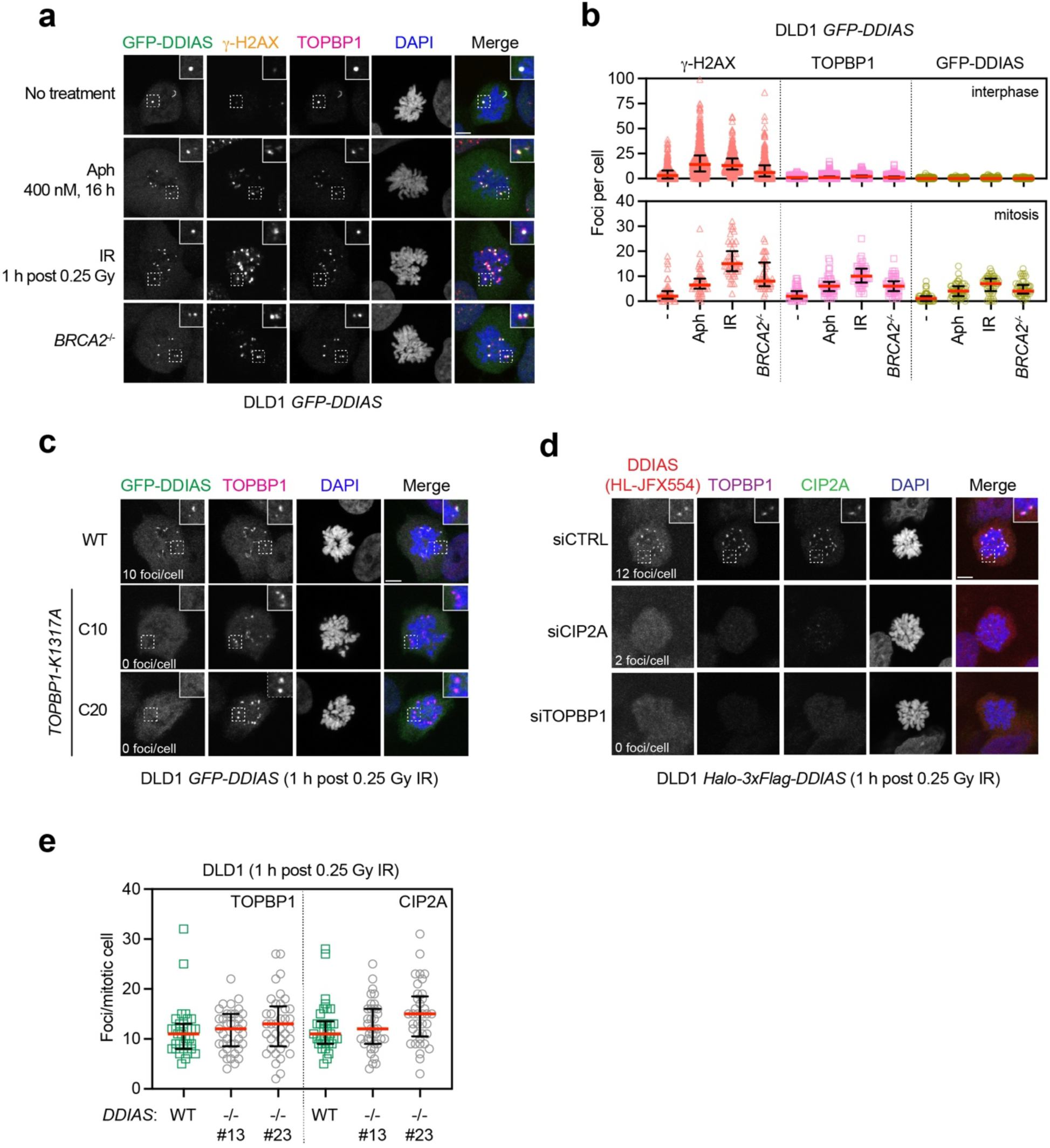
DDIAS localizes to mitotic DNA damage sites. (**a**) Representative micrographs showing the localization of GFP-DDIAS on DNA lesions induced by the indicated conditions in mitotic DLD1 cells. (**b**) Quantification of GFP-DDIAS, γ-H2AX and TOPBP1 foci in mitotic and interphase cells in **a**. Bars indicate the mean ± SD, with n=3 independent experiments. (**c**) Representative images of GFP-DDIAS foci in mitotic WT and TOPBP1 K1317A-mutated DLD1 cells after IR treatment. Average numbers of GFP-DDIAS foci from 3 independent experiments are indicated. Quantification of GFP-DDIAS and TOPBP1 foci in individual mitotic cells is presented in Fig. S5d. (**d**) Immunofluorescence analysis of endogenous DDIAS localization in mitotic DLD1 Halo-DDIAS cells treated with siRNAs against CIP2A or TOPBP1 followed by IR exposure. Halo-DDIAS was detected by Janelia Fluor JFX554 HaloTag Ligand. Median numbers of GFP-DDIAS foci from 3 independent experiments are indicated. Quantification of DDIAS, CIP2A and TOPBP1 foci in individual mitotic cells is presented in Fig. S5e. (**e**) Immunofluorescence analysis of TOPBP1 and CIP2A foci in mitotic DLD1 cells with indicated genetic background following IR treatment. Bars indicate the mean ± SD. n=3 independent experiments. Scale bar, 5 μm.

Next, we inserted the HaloTag-coding sequence^42^ flanked by HiBiT^43^ and 3xFlag at the chromosomal *DDIAS* locus in DLD1 cells to produce endogenously tagged DDIAS (referred hereafter as *Halo-DDIAS*; Fig S4c,d), as we have yet to find an antibody that can detect endogenous levels of DDIAS. Halo-DDIAS is recruited to mitotic DNA damage foci that also contain CIP2A and TOPBP1 in a manner that depends on both proteins (Fig 3d and Fig S5f). Since the loss of DDIAS did not impact the localization of CIP2A or TOPBP1 at mitotic DNA lesions (Fig 3e and Fig S5g), we conclude that DDIAS is a bona fide mitotic DNA damage response protein that is co-recruited with the CIP2A-TOPBP1 complex in a manner that depends on phosphopeptide-binding by the TOPBP1 BRCT7/8 domain.

### DDIAS phospho-dependent interaction with TOPBP1

We next sought to determine whether DDIAS interacts directly with the TOPBP1 BRCT7/8 domain. As a first step, we carried out structure-function studies to determine the minimal fragment of DDIAS that is competent for accumulation at sites of mitotic DNA damage. These efforts converged on a C-terminal fragment (DDIAS [651-998]; Fig S6a) able to form mitotic foci as efficiently as full-length DDIAS (Fig S6a-d). Interestingly, this region did not encompass the OB-fold domain (Fig S6a), suggesting that its putative nucleic acid-binding activity was not involved in its DNA damage localization. To further narrow down the key residues involved in the DDIAS-TOPBP1 interaction, we employed AlphaFold3^44^ to build a model of phosphorylated DDIAS bound to the BRCT7/8 domains of TOPBP1, which revealed the DDIAS sequence surrounding residue S676 as the top-ranked potential BRCT7/8-binding site (Fig 4a). The sequence of the S676-containing epitope is highly similar to that of the BRCT7/8-binding segment on BRIP1, anchored by T1133, whose structure was determined by X-ray crystallography^45^ (Fig 4b). Generation of a DDIAS S676A mutant reduced, but did not completely abolish, the mitotic DNA damage localization of DDIAS (Fig 4c and Fig S6e,f).

**Figure 4.**
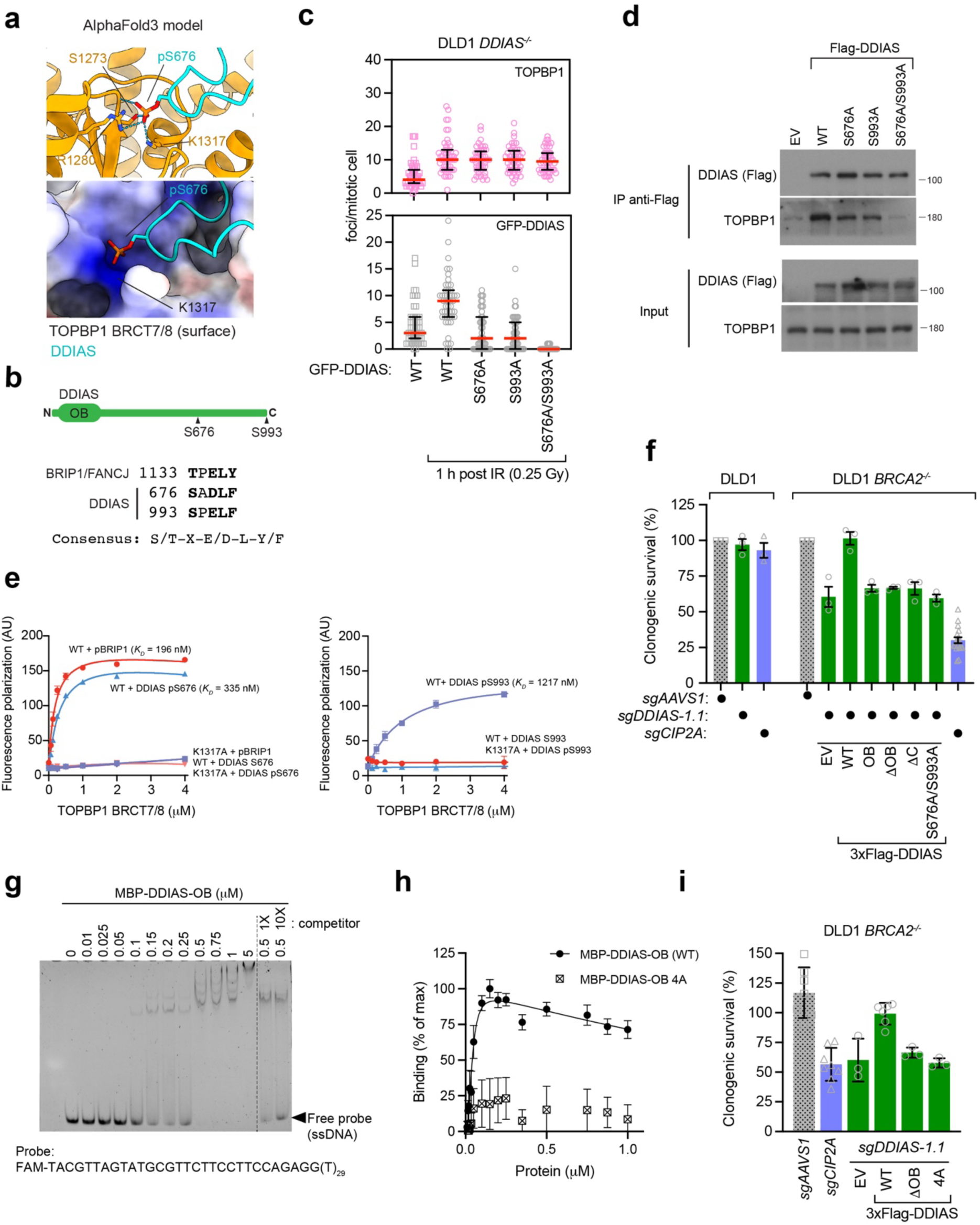
Characterization of the DDIAS-TOPBP1 interaction. (**a)** AlphaFold3 prediction of TOPBP1 BRCT7/8 in complex with DDIAS (631-720) containing phosphorylated S676. (**b)** The sequences of TOPBP1 BRCT7/8-binding segments on DDIAS and BRIP1. (**c)** Immunofluorescence analysis of TOPBP1 and GFP-DDIAS foci in mitotic DLD1 cells expressing the indicated variants of GFP-DDIAS with or without IR treatment. Bars indicate the mean ± SD. n=3 independent experiments. (**d)** Immunoblots showing the co-immunoprecipitation of TOPBP1 and the indicated 3xFlag-DDIAS variants in HEK293 cells. n=2 independent experiments. (**e)** Fluorescence polarization assay measuring the binding of indicated FITC-labeled peptides to recombinant WT or K1317A-mutated BRCT7/8 of TOPBP1. Bars indicate the mean ± SD. n=3 independent experiments. (**f)** Clonogenic survival of DLD1 WT and *BRCA2*^-/-^ cells expressing the indicated sgRNA-resistant *DDIAS* variants or EV (empty vector), followed by depletion of endogenous DDIAS or CIP2A by sgRNAs. Bars indicate the mean ± SEM. nζ 3 independent experiments. (**g)** Electrophoretic mobility shift assay (EMSA) using recombinant MBP-DDIAS-OB and fluorescently labeled single-stranded (ss) DNA with or without unlabeled competitor. Representative gel image shown from n=3 independent experiments. (**h)** Time-resolved fluorescence energy transfer (TR-FRET) assay using fluorescently labeled ssDNA and increasing concentration of recombinant WT or 4A mutated MBP-DDIAS-OB. Signal was normalized to maximal binding activity by WT. Bars indicate the mean ± SEM. N=3 independent experiments. (**i**) Clonogenic survival of DLD1 *BRCA2*^-/-^ cells expressing the indicated sgRNA-resistant *DDIAS* variants or EV (empty vector), followed by depletion of endogenous DDIAS or CIP2A by sgRNAs. Bars indicate the mean ± SEM. nζ 3 independent experiments.

We therefore searched for additional phosphorylatable sites in DDIAS that could mediate an interaction with TOPBP1 in parallel to S676 and identified S993, at the extreme C-terminus of DDIAS (Fig 4b). The double S676A/S993A mutation completely abolished DDIAS foci in mitosis without altering TOPBP1 localization (Fig 4c and Fig S6e,f). Finally, DDIAS S676A/S993A is unable to interact with TOPBP1 as it does not co-immunoprecipitate TOPBP1 from mitotic whole cell extracts (Fig 4d). These data suggest that DDIAS phosphorylated at the S676 and S993 sites may be recognized by TOPBP1.

To assess whether DDIAS S676 and S993 phosphorylation can promote a direct interaction with the TOPBP1 BRCT7/8 domain, we synthesized phosphopeptides encompassing either site (DDIAS pS676 and pS993) and tested them for interaction with recombinant BRCT7/8 in fluorescence polarization assays. As a positive control, we used a synthetic BRIP1-derived peptide (pBRIP1, encompassing the phosphorylated T1113 residue). We found that both DDIAS-derived phosphopeptides, but not their unphosphorylated counterparts, interact with recombinant BRCT7/8 in a K1317 residue-dependent manner with *K_D_* values (0.3 μM and 1.2 μM for pS676 and pS993, respectively) similar to that of the pBRIP1-BRCT7/8 interaction (0.2 μM; Fig 4e). We conclude that DDIAS is recruited to mitotic DNA lesions via direct phospho-dependent interactions with the TOPBP1 BRCT7/8 domain. These studies also enable us to derive a new consensus BRCT7/8-binding sequence: pS/pT-X-D/E-L-Y/F, where pS/pT are phosphorylated Ser/Thr residues and X represents any amino acid residue (Fig 4b).

### The DDIAS-TOPBP1 interaction is essential in *BRCA2^-/-^* cells

Next, we mapped the DDIAS regions that promote cell viability under conditions of HR deficiency. To do so, we generated DDIAS lentiviral expression vectors that are resistant to the *DDIAS*-targeting sgRNA *sgDDIAS1.1*. These vectors express Flag-tagged DDIAS, or versions that delete the N-terminal OB-fold (DDIAS-ΔOB, corresponding to residues 198-998), that delete the C-terminal region (DDIAS-ΔC, corresponding to residues 2-800), that express solely the OB-fold domain (DDIAS-OB, corresponding to residues 2-197) or that encode the DDIAS S676A/S993A mutant (Fig S6g). These lentiviruses, alongside an empty virus (EV) control, were introduced in DLD1 *BRCA2^-/-^*cells. These cells were then infected with a second lentivirus that expressed sgRNAs targeting *AAVS1*, *CIP2A* or *DDIAS*. As controls, we also infected wild type DLD1 with the sgRNA-expressing viruses. As expected, CIP2A and DDIAS depletion caused loss of clonogenic survival only in *BRCA2^-/-^* cells, and the loss of viability imparted by *sgDDIAS1.1* was completely suppressed by the exogenous expression of the sgRNA-resistant DDIAS transgene (Fig 4f and Fig S6h). Expression of DDIAS-ΔC or DDIAS S676A/S993A were unable to suppress the *DDIAS/BRCA2* synthetic lethality, and neither did DDIAS-ΔOB or DDIAS-OB (Fig 4f and Fig S6h). These data suggest that formation of the CIP2A-TOPBP1-DDIAS complex promotes the viability of *BRCA2^-/-^* cells and that the OB-fold containing region of DDIAS may also contribute to the fitness of HR-deficient cells.

### The OB-fold of DDIAS binds to ssDNA

The importance of the N-terminal region for DDIAS function suggests that the OB-fold mediates a critical activity of the CIP2A-TOPBP1 complex. Given that OB-folds are often nucleic acid binding-domains^41^, we tested the capacity of a recombinant DDIAS OB-fold, produced in *Escherichia coli* as an MBP fusion protein (Fig S7a), to bind to DNA. We first used electrophoretic mobility shift assays (EMSA) with fluorescently labeled single-stranded (ss) or double-stranded (ds) DNA probes and found that DDIAS OB-fold binds to ssDNA preferentially (Fig 4g and S7b). Dose-dependent ssDNA-binding by the DDIAS OB-fold could also be detected using time-resolved fluorescence energy transfer (TR-FRET) assays, which are conducted in solution (Fig S7c). The ssDNA-binding signal observed by EMSA and TR-FRET could be competed with excess cold ssDNA probe (Fig 4g and S7d), suggesting that the binding can be saturated and is thus specific.

To assess the importance of ssDNA-binding for DDIAS function, we searched for DNA-binding mutants. Given that DNA binding often involves interactions between positively charged amino acid residues and the phosphodiester backbone, we generated an OB-fold mutant that harbors the quadruple R3A, R31A, R56A and K58A mutation (referred to as “4A”) that target basic residues at the predicted OB-fold surface (Fig S7e). The recombinant OB-fold 4A mutant was highly defective in ssDNA-binding, as measured by TR-FRET (Fig 4h). We then generated an sgRNA-resistant lentivirus expressing the corresponding Flag-DDIAS 4A mutant and found that this mutant was unable to rescue the viability of DLD1 *BRCA2^-/-^* cells infected with a DDIAS-targeting sgRNA (Fig 4i and S7f,g) despite being able to localize to mitotic DNA lesions (Fig S7h). These data indicate that DNA binding by DDIAS is central to the function of the CIP2A-TOPBP1-DDIAS complex in mediating genome integrity but is dispensable for its localization to mitotic DNA damage sites.

### DDIAS buffers defective fill-in synthesis

Consistent with the observation that TOPBP1-K1317A does not impair micronuclear chromatin fragment clustering in mitosis, we found that *DDIAS^-/-^*cells cluster fragments arising from micronuclear chromosome shattering much more efficiently than *CIP2A^-/-^* cells, suggesting that DDIAS does not act primarily to tether broken DNA ends (Fig S8a). To gain insight into the mechanism by which DDIAS acts to promote genome integrity, we undertook a genome-scale synthetic lethality screen to identify genes whose inactivation have impaired viability in the absence of DDIAS. We profiled parental DLD1 and DLD1 *DDIAS^-/-^* cells with the lentiviral TKOv3 sgRNA library^46^ and computed gene-level essentiality scores (Bayes Factor, or BF) using BAGEL2^47^ (Fig 5a and Table S4). We identified 73 genes whose sgRNAs caused selective loss of fitness in *DDIAS^-/-^* cells (ΔBF_(KO-WT)_>10 with BF_WT_ <20 and BF_KO_>10; Fig 5a and Table S4). Analysis of this gene list for Gene Ontology (GO) Biological Process terms found enrichment in multiple terms associated with DNA repair and mitotic cell division (Fig S8b) consistent with a role for DDIAS in the mitotic DNA damage response.

**Figure 5.**
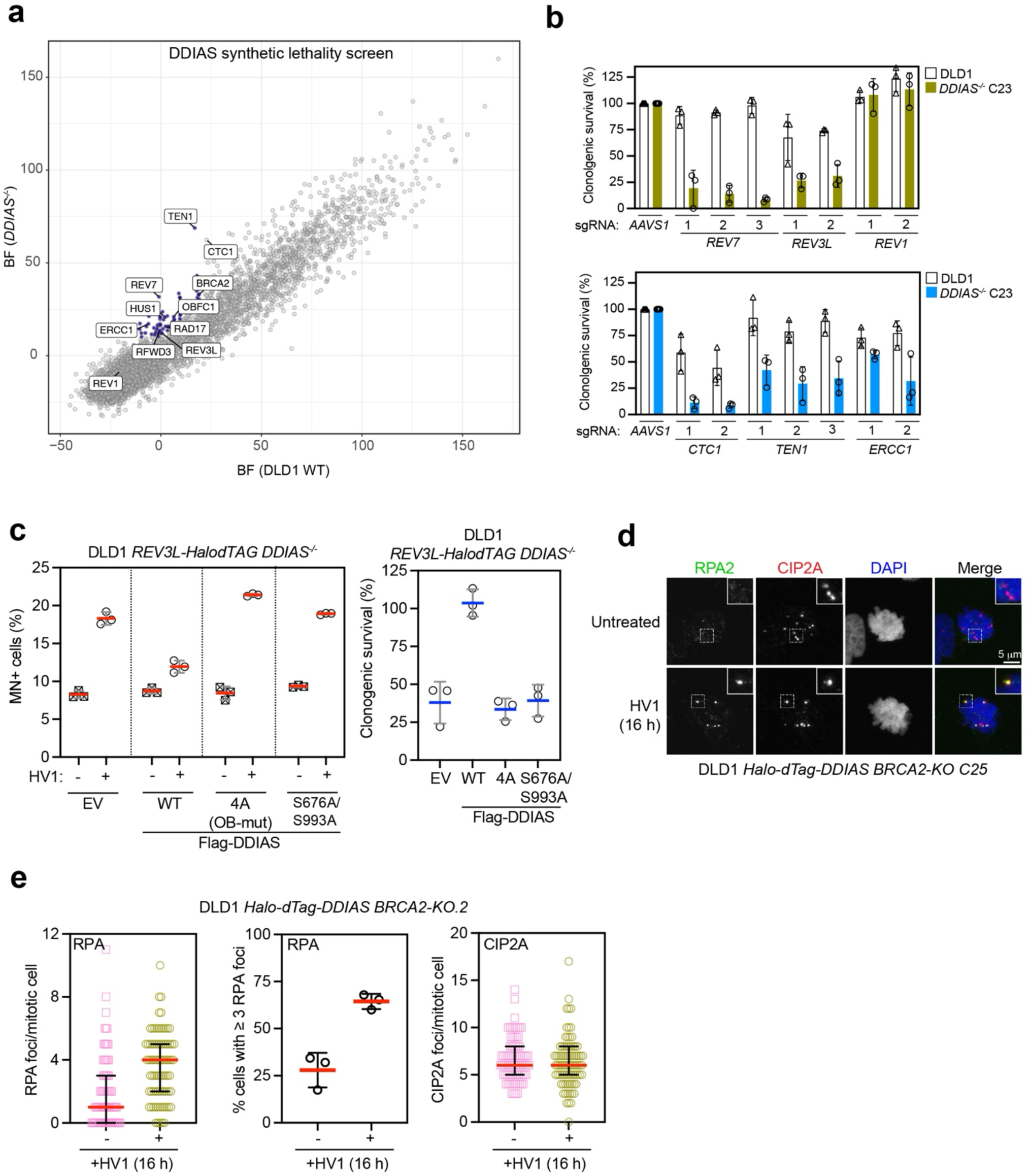
DDIAS counteracts fill-in synthesis defects and suppresses mitotic ssDNA. (**a)** Scatter plot of Bayes factor (BF) values derived from BAGEL2 for the DDIAS isogenic synthetic lethal screen performed in DLD1 cells. (**b**) Clonogenic survival of DLD1 WT and *DDIAS*^-/-^ cells expressing the indicated sgRNAs. Bars indicate the mean ± SD. n=3 independent experiments. (**c**) Quantification of micronuclei and clonogenic survival in DLD1 *REV3L-HalodTAG DDIAS^-/-^* cells expressing the indicated variant of DDIAS with or without REV3L depletion. HV1 presents HaloPROTAC3 (1 μM) + dTAG-v1 (1 μM). Clonogenic survival data was normalized to untreated condition. Bars indicate the mean ± SD. n=3 independent experiments. (**d**) Immunofluorescence analysis of RPA and CIP2A foci in mitotic DLD1 *HalodTAG-DDIAS BRCA2*^-/-^ cells with or without DDIAS depletion. (**e**) Quantification of the RPA and CIP2A foci from the experiment shown in **d**. Bars indicate the mean ± SD. n=3 independent experiments. Scale bar, 5 μm.

Among the top genes promoting viability of *DDIAS^-/-^* cells were *BRCA2*, as expected, as well as *REV3L* and *REV7* (also known as *MAD2L2*), which code for two components of Pol(^48^ as well as genes encoding components of the CST complex (CTC1, TEN1 and OBFC1/STN1; Fig 5a and Table S4). Given that Pol(and CST promote fill-in synthesis (the latter via Polα-primase^49^), the genetic interaction profile hints at DDIAS being important for cell fitness when fill-in DNA synthesis is compromised. Using independent sgRNAs, we ascertained that depletion of REV3L, REV7, CTC1 and TEN1 caused loss of clonogenic survival in *DDIAS^-/-^* cells but not in their parental counterpart (Fig 5b and S8c,d). Interestingly, we found that loss of REV1, involved in the translesion DNA synthesis (TLS) function of Pol(, is viable in *DDIAS^-/-^* cells, implicating a TLS-independent function of Pol(in the *DDIAS-REV3L/REV7* synthetic lethality (Fig 5b and S8c,d). We also tested whether *ERCC1*-targeting sgRNAs impaired viability of *DDIAS^-/-^* cells given the recent links between ERCC1-XPF and the CIP2A pathway^21,50^, but the results were inconclusive despite efficient ERCC1 depletion (Fig 5b and S8c,d) The combined loss of DDIAS and either REV3L or REV7 caused a marked increase in micronucleation whereas depletion of CST-coding genes in *DDIAS^-/-^* cells had more modest impact (Fig S8e). Finally, we observed that *DDIAS^-/-^* cells are viable in the absence of the SHLD2 and SHLD3 subunits of the shieldin, a DNA repair complex that contains REV7 and that interacts with CST^51–55^. The lack of genetic interaction between *DDIAS* and shieldin-coding genes suggests that it is the role of REV7 within Pol(that is essential when DDIAS is absent (Fig S8f). Together, these results suggest that DDIAS may be particularly important in counteracting DNA lesions that requires fill-in synthesis.

Given that Pol(loss caused the most pronounced phenotypes in the absence of DDIAS, we focused on this genetic interaction. To avoid complications associated with the impaired cell fitness caused by long-term REV3L loss, we constructed a DLD1 derivative in which we endogenously tagged REV3L with both HaloTag and dTAG (*REV3L-HaloTag-dTAG-3xFlag-HiBiT*, shortened to *REV3L-HalodTag*; Fig S9a). This cell line allowed for rapid depletion of REV3L within ∼1 h with a combination of HaloPROTAC3 and dTAG-1 (condition HV1; Fig S9b). REV3L-depleted cells accumulated TOPBP1 and γ-H2AX mitotic foci 24 h after HV1 addition, indicating that Pol(loss causes DNA lesions that are transmitted into mitosis (Fig S9c). Generation of a *DDIAS^-/-^* derivative in the *REV3L-HalodTAG* cell line allowed us to confirm that loss of Pol(caused micronucleation and lethality in the absence of DDIAS, and importantly, we also ascertained that REV3L depletion impaired viability of *CIP2A^-/-^*cells as well (Fig 9Sd). Consistent with this result, we found that Cas9-mediated inactivation of REV3L and REV7 reduced the clonogenic survival of *CIP2A^-/-^* and *TOPBP1-K1317A* cells (Fig S9e). In all cases, impaired cell viability was accompanied by increased genomic instability as measured by micronucleation (Fig S9d,f). To further assess whether Pol(loss caused a reliance on the mitotic CIP2A-TOPBP1-DDIAS pathway, we expressed DDIAS WT or the DDIAS-S676A/S993 and -4A mutants in *REV3L-HalodTAG DDIAS^-/-^* cells and found that disruption of the interaction with TOPBP1 or ssDNA caused loss of viability and higher levels of micronucleation in cells depleted of REV3L (Fig 5c and S9g). Together, these data indicate that cells lacking fill-in synthesis by Pol(rely on the CIP2A-TOPBP1-DDIAS complex for viability (and genome stability) in a manner that relies on the ssDNA-binding activity of DDIAS.

### DDIAS promotes resolution of ssDNA-containing mitotic DNA lesions

The most parsimonious model to explain how a mitotic ssDNA-binding protein may promote genome integrity when fill-in synthesis is disabled, is that DDIAS promotes the resolution of ssDNA-containing DNA lesions. To explore this model, we assessed whether DDIAS depletion in the *HalodTAG-DDIAS BRCA2-KO* cell line increases mitotic ssDNA lesions detected with RPA immunofluorescence as proxy. We previously reported that *BRCA2^-/-^* cells have 1-2 RPA foci in early mitotic cells that likely originate from replication-associated DNA lesions^7^. Depletion of DDIAS caused an increase in RPA foci in *BRCA2-KO* cells (Fig 5d,e), but not an increase in CIP2A foci, suggesting that DDIAS acts on ssDNA-containing DNA lesions downstream of CIP2A. Consistent with this idea, most RPA foci co-localized with CIP2A (Fig 5d and S10a).

Next, we exogenously expressed DDIAS, DDIAS-4A or the S676A/S993A variant in *DDIAS-HalodTAG BRCA2-KO* cells. We found that while exogenous expression of DDIAS WT rescued RPA foci upon degradation of HalodTAG-DDIAS, neither DDIAS-4A nor -S676A/S993A did so (Fig S10b). These results suggest that the CIP2A-TOPBP1-DDIAS complex suppresses mitotic ssDNA-containing DNA lesions in a manner that depends on the DDIAS ssDNA-binding activity.

Given the genetic interaction between Pol(and both CIP2A and DDIAS, we similarly stained mitotic cells for RPA foci upon depletion of REV3L in the *REV3L-HalodTAG* cell line and in its *CIP2A^-/-^* and *DDIAS*^-/-^ counterparts. We found that in cells with functional CIP2A and DDIAS (i.e. the WT condition), depletion of REV3L had a negligible impact on a low basal level of mitotic RPA foci (Fig S10c). However, we could detect a small but reproducible increase in RPA foci upon Pol(depletion in both *CIP2A^-/-^* and *DDIAS^-/-^* cells (Fig S10c). These data raise the possibility that a core function of the CIP2A-TOPBP1-DDIAS complex is to act on mitotic ssDNA-containing lesions.

## DISCUSSION

The set of processes that manage mitotic DNA lesions constitutes a genome “rescue” system that minimizes the negative impact of chromosome breakage in mitosis, and that participates in the orderly disassembly of DNA replication and recombination intermediates on mitotic chromosomes. The importance of these pathways is illustrated by the fact that the CIP2A-dependent mitotic DNA damage response is broadly essential in HR-deficient cells^7^ and promotes resistance to many genotoxic agents^56^. However, the origin of the mitotic DNA lesions that compromise genetic integrity and viability in BRCA-deficient cells (and in HR-proficient cells) is not well understood. Here, we report that the ssDNA-binding protein DDIAS forms a mitotic DNA damage response complex with CIP2A and TOPBP1 that promotes genome stability and viability of HR-deficient cells. Characterization of DDIAS leads us to propose that mitotic ssDNA represents a liability for chromosome stability in mitotic cells.

Exactly why the CIP2A pathway requires a ssDNA-binding protein was at first puzzling, but our data paints a picture in which CIP2A-TOPBP1 does not anchor a singular end-tethering pathway during mitosis but rather organizes distinct mitotic processes that promote genomic integrity. This is illustrated by the observation that unlike CIP2A, both the TOPBP1 BRCT7/8 module and DDIAS are largely dispensable for the clustering of shattered micronuclear chromosomes, but both are required for chromosomal stability in HR-deficient cells. These observations, along with our structure-function studies reveal that CIP2A-TOPBP1 forms a subcomplex with DDIAS, which is critical to promote the resolution of mitotic ssDNA-containing lesions that likely arise due to defective DNA replication, in a manner that involves the ssDNA-binding OB-fold of DDIAS. This is evidenced by the increase in mitotic RPA foci when BRCA2 and REV3L deficiency is combined with DDIAS loss. However, we note that RPA mitotic foci do not perfectly correlate with micronucleation and lethality, raising the possibility that RPA-bound ssDNA may not be the sole type of DNA lesion that DDIAS helps resolve. The formation and extension of replicative ssDNA gaps during interphase is increasingly recognized as a pathological feature that endangers genomic integrity and chemotherapeutic responses^57–59^. However, it remains unclear why gaps produced during DNA replication or upon conditions of HR deficiency are such a threat. Our study suggests that mitotic ssDNA lesions, generated either during mitosis or transmitted from interphase, could represent a source of mitotic chromosome breakage that is counteracted by CIP2A-TOPBP1-DDIAS. This idea is appealing since forces in the range of 1-2 nN are predicted to be sufficient to break ssDNA^5,60^, which is in the same range as the forces generated by the mitotic spindle^61^. Therefore, mitotic ssDNA has potential to generate chromosome breakage by both mechanical and enzymatic means, potentially causing acentric fragment mis-segregation, micronucleation and chromothriptic-like chromosome rearrangements. This model, while attractive, is unlikely to fully explain how interphase replicative ssDNA gaps compromise genomic integrity, but it provides avenues of investigation that will form the basis of future studies.

While we present a simplified model of CIP2A-TOPBP1-DDIAS function, the reality is clearly more complex. Indeed, the phenotypes of the TOPBP1-K1317A mutant are not identical to those of *DDIAS*^-/-^ cells, which is not surprising given that TOPBP1 interacts with multiple proteins involved in genome maintenance via the BRCT7/8 module. Candidate proteins that may make additional contributions to the CIP2A-TOPBP1 pathway via the BRCT7/8 module include Polθ^23^, TTF2 as well as PLK1. In addition to these interactions, CIP2A-TOPBP1 also interacts with the SLX4-anchored SMX nuclease complex^62^ to promote CIP2A-dependent processes^21,50^. CIP2A-TOPBP1 is thus emerging as an organizing platform for mitotic chromosome integrity and the identification of DDIAS and its role in suppressing mitotic ssDNA highlights the multifaceted role of this pathway during cell division.

## Acknowledgments

We thank Rachel Szilard to help with editing this manuscript; Peter Ly and Yu-Fen Lin for the CEN-SELECT parental and *CIP2A^-/-^* cell lines, as well as useful information on the micronuclear chromosome tethering assay; Xialu Li for the REV3L antibody; Dr. Payman Samavarchi-Tehrani and Hala Abdouni for help with designing, cloning, and testing the split-TurboID constructs, and Andy Blackford for collegial discussions on DDIAS. The authors wish to thank Annie Bang, Kin Chan, Monica Hasegan, Johnny Tkach, Cassandro Wong of the Network Biology Collaborative Centre facilities (RRIDs: SCR_025375, SCR_025385 and SCR_025389) at the Lunenfeld-Tanenbaum Research Institute for their expert contribution to this work. The NBCC is supported by the Canada Foundation for Innovation and the Ontario Government. We dedicate this work to the memory of Dr. Silvia Emma Rossi, who first noted the DDIAS-BRCA genetic interaction. YX was supported by CHIR Postdoctoral Fellowship and a Hold’em for Life Oncology Fellowship. Work in the FS and ACG labs was supported by Terry Fox New Frontiers Program Project Grant. Work in the DD lab was supported by grants from the Canadian Institutes for Health Research (CIHR grant PJT 180438) and with support from the Dani Reiss Innovation Fund for Healthy Ageing.

## Author Contributions

Conceptualization, Investigation, Validation, Writing and Visualization, Y.X.; Conceptualization, Investigation, Validation, Writing and Visualization, F.R.; Investigation, D.Y.L.M.; Resources, Methodology, K.T.A.; Investigation, Z.-Y.L.; Investigation, Visualization, D.S.; Formal Analysis, Data Curation, Visualization, L.H.; Validation, C.B.; Supervision, F.S.; Resources, Methodology, Supervision, A.-C.G.; Conceptualization, Supervision, Writing and Funding Acquisition, D.D.

## Methods

### Cell culture

The RPE1-hTERT and 293T cell lines were grown in Dulbecco’s modified Eagle’s medium (DMEM) supplemented with 10% FBS (Wisent catalog no. 080150), 1% GlutaMax (Gibco, catalog no. 35050-061) and 1% penicillin-streptomycin (Wisent #450-201-EL). Parental DLD-1 cells (Horizon Discovery, catalog no. HD 105-007) and genetically engineered derivatives were maintained in RPMI-1640 medium (American Type Culture Collection (ATCC), catalog no. 30-2001) supplemented with 10% FBS, 1% GlutaMax and 1% penicillin-streptomycin. All cell lines were cultured at 37 °C under 5% CO_2_ and atmospheric oxygen except *BRCA2^-/-^* cells which were maintained in an incubator with 3.5% oxygen. Cell lines were routinely monitored to ensure the absence of mycoplasma contamination. Parental cell lines have been authenticated by short tandem repeat analysis.

### Cell line engineering

The conditional silencing system for TOPBP1was generated by adopting a previously described method^63^, where TOPBP1 expression is modulated by the combination of a tetracycline-controlled (Tet-Off) promoter at the transcriptional level and degron-mediated regulation for protein degradation. Specifically, HA-TOPBP1 harboring a sgRNA-resistant silent mutation was cloned into plasmid pUHD-SB-mAID/Hyg (Addgene: #171679), and the F74G mutation of AID2^64^ was introduced into for *OsTIR1* in plasmid pSBbi-TIR1-tTA/Pur (Addgene: #171683). These two plasmids, together with the SB transposase plasmid (pCMV(CAT)T7-SB100; Addgene, #34879), were transfected into DLD1 cells with Lipofectamine 3000 Transfection Reagent (Invitrogen, # L3000001). Two days after transfection, cells were selected with hygromycin and puromycin for two weeks. Endogenous *TOPBP1* in the recovered cells was knocked out as described below.

To introduce the K1317A mutation on endogenous *TOPBP1*, a homology-directed repair (HDR) donor oligo and an associated sgRNA were designed with the Alt-R HDR Design Tool provided by Integrated DNA Technologies, Inc. Following the manufacture’s instruction, the HDR donor oligo and CRISPR-Cas9 ribonucleoprotein complex were delivered into DLD1 cells by using electroporation with Lonza 4D Nucleofector Core Unit and SF Cell Line 4D-Nucleofector X Kit S (Lonza, # V4XC-2032). Two days after electroporation, 200 cells were seeded in a 15-cm dish for clonal growth. After ∼10 days, genomic DNA from expanded cell clones were isolated with DNeasy Blood and Tissue Kit (Qiagen # 410017809), and the genomic region surrounding TOPBP1 K1317 in the clones was amplified by PCR using KAPA HiFi HotStart ReadyMix (Kapa Biosystems, KK2602). PCR products were sequenced to verify the K1317A mutation.

HDR donor template:

5’-TGA TCC CAC CTG TAC ACA CAT TGT TGT GGG ACA TCC ACT TCG AAA CGA GGC GTA TTT AGC CTC AGT GGC AGC TGG GAA GTG GGTG CTT CAT-3’ sgRNA: AAATACTTCTCGTTTCGAAG Endogenous tagging of *TOPBP1*, *DDIAS* and *REV3L* with HaloTag and/or dTAG was carried out as previously described with minor modifications^11^. sgRNA targeting the desired genomic regions (Fig S3a, S4c, S9a) were designed by the Benchling CRISPR Guide RNA design tool and cloned into pSpCas9(BB)-2A-GFP (PX458) (Addgene # 48138). The left and right homology arms (approximately 500 bp) were synthesized by Twist Bioscience and cloned into pBluescript II KS (-) (Stratagene). For each tagging experiment, sgRNA and template plasmids were transfected together into DLD1 or DLD1-ChY cells with Lipofectamine 3000 transfection reagent. GFP expressing single cells were sorted into 96-well plates for colony growth two days post transfection. After ∼10 d, genomic DNA from expanded cell clones were isolated and analyzed by PCR to identify clones that were correctly edited. Finally, PCR products were sequenced for further verification.

To knock out *CIP2A*, *DDIAS* and *BRCA2* in DLD1 derivatives, CRISPR-Cas9 ribonucleoprotein complexes with associated sgRNA were delivered into cells by electroporation as described above. Two days post-electroporation, single cells were sorted into 96-well plates for colony growth. After ∼10 d, genomic DNA from expanded cell clones was isolated and the targeted genomic regions were amplified by PCR. PCR products were sequenced, and genomic editing was evaluated by TIDE^65^ or the Synthego ICE analysis tool. The sgRNA sequences and PCR primers used for clone verification are included in Table S3.

### Plasmids

For CRISPR-Cas9 genome editing, sgRNAs were cloned either into lentiCRISPRv2 or in lentiguide-NLS-GFP as described^66^. The sgRNA sequences used in this study are included in Table S3. pCIB-HA2-TOPBP1 FL (sg10R) was from a previous study^7^. DDIAS coding sequences with GFP, Flag or HA tags were cloned into pHIV or pCIB^7^ plasmids using Gibson assembly, which led to the generation of pHIV-GFP-DDIAS, pHIV-3xFlag-DDIAS, pCIB-3xFlag-DDIAS, pCIB-GFP-DDIAS and pCIB-2xHA-DDIAS. Unless stated otherwise, both TOPBP1 and DDIAS mutant plasmids were generated by site-directed mutagenesis using Q5 High-Fidelity 2X Master Mix (NEB, # M0492S). Plasmid sequences were verified by Plasmidsaurus Inc or the TCAG DNA sequencing facility at the Hospital for Sick Children. Plasmids used for genome editing, Split-TurboID and protein production are described in related sections.

### siRNAs

siGENOME non-targeting control siRNA Pool #2 (Dharmacon, # D-001206-14-20), ON-TARGETplus pool KIAA1524 (CIP2A) siRNA (Dharmacon, # L-014135-01-0005), and siGENOME SMARTpool TOPBP1 siRNA (Dharmacon, # M-012358-01-0005) were used. Transfection was performed using Lipofectamine RNAiMAX (Thermo Fisher Scientific, #13778150) by following the manufacture’s instruction.

### Fine chemicals and antibiotics

Nocodazole (Sigma, #M-1404), aphidicolin (Focus Biochemicals, #10-2058), olaparib (SelleckChem, # S1060), doxycycline hyclate (Sigma, #D5207), AquaShield-1 (CheminPharma, #AS1-0005), indole-3-acetic acid (Sigma, #I5148), 5-Ph-IAA (MedChem Express, #HY-134653), dTAG-13 (Tocris Bioscience, #6605), dTAGV-1 (Tocris Bioscience, #6914), HaloPROTAC3 (Glixx Laboratories, #GLXC-15215), Janelia Fluor JFX554 HaloTag Ligand (Promega, #HT1030), puromycin dihydrochloride (Life Technologies, #A1113802), blasticidin S HCl (Life Technologies, #R21001), geneticin (G418) (Gibco, #10131-035), hygromycin (Millipore Sigma, #10843555001), nourseothricin sulfate (Gold Biotech, # N-500-5)

### Antibodies

The antibodies listed below were used for immunoblotting (IB) or immunofluorescence (IF). Primary antibodies: mouse anti-CIP2A (clone 2G10-3B5; Santa Cruz sc80659, 1:500 IF, 1:1000 IB), rabbit anti-CIP2A (Novus Biologicals # NBP2-48710, 1:1000 IF), rabbit anti-phospho-Histone H2A.X (Ser139) (Cell Signalling Technologies #2577, 1:1000 IF), mouse anti-phospho-Histone H2A.X (Ser139) (clone JBW301; Millipore Sigma #05-636, 1:5000 IF), mouse anti-FLAG M2 (Sigma G1804, IB 1:1000), rat anti-FLAG (BioLegend #637301, 1:1000 IF), rabbit anti-TOPBP1 (Abcam ab2402, 1:2000 IF, 1:5000 IB), mouse anti-alpha-tubulin (Cell Signalling Technologies #3873, 1:10,000 IB), mouse anti-HA.11 (clone 16B12, BioLegend # 901502, 1:200 IF), rabbit anti-HA (Cell Signalling Technologies #3724, 1:1000 IB), rabbit anti-REV3L (a gift from Xialu Li, 1:1000 IB), mouse anti-ERCC1 (Santa Cruz sc-17809, 1:200 IB), rabbit anti-REV7(Abcam, ab180579, 1:1000 IB), mouse anti-REV1 (Santa Cruz sc-393022, 1:500 IB), rabbit anti-GFP (gift from Laurence Pelletier, 1:5000 IB), rabbit anti-Myc (Cell Signalling Technologies #2278, 1:5000 IB), rat anti-RPA2 (Cell Signalling Technologies #2208, 1:10,000 IF), human anti-centromere protein (Antibodies Incorporated, #15-235 1:1000 IF), rabbit anti-BRCA2 (Abcam, ab27976, 1:1000 IB), Rabbit anti-KAP1 (Bethyl, #A300-274A, 1:10,000 IB).

Secondary antibodies for immunoblotting: IRDye 680RD goat anti-mouse IgG (Li-Cor, #926-68070, 1:5000), IRDye 800CW Streptavidin (Li-Cor, #926-32230, 1:5000), HRP-conjugated bovine anti-goat IgG (Jackson ImmunoResearch, #805-035-180, 1:5000), HRP-conjugated goat anti-rabbit IgG (Jackson ImmunoResearch, #111-035-144, 1:5000), and HRP-conjugated goat anti-mouse IgG (Jackson ImmunoResearch, #115-035-003, 1:5000).

Secondary antibodies for immunofluorescence: AlexaFluor 488-goat anti-mouse IgG (Thermo Fisher Scientific, #A11029, 1:1000), AlexaFluor 555-goat anti-mouse IgG (Thermo Fisher Scientific, #A21424, 1:2000), AlexaFluor 647-goat anti mouse IgG (ThermoFisher Scientific, #A21236, 1:2000), AlexaFluor 647-goat anti-rabbit IgG (ThermoFisher Scientific, #A21244, 1:2000 IF), AlexaFluor 488-goat anti-rabbit IgG (Thermo Fisher Scientific, #A11034, 1:2000), AlexaFluor 555-goat anti-rabbit IgG (Thermo Fisher Scientific, #A21428, 1:2000), AlexaFluor 488-goat anti-rat IgG (Thermo Fisher Scientific, #A11006, 1:1000), AlexaFluor 488-goat anti-human IgG (Thermo Fisher Scientific, #A11013, 1:1000)

### Lentiviral transduction

To produce lentiviral particles, 293T cells were seeded in 6-well plate with 2x10^6^ cells per well. Packaging plasmids (1 μg pVSVg, 0.5 μg pMDLg/pRRE and 0.5 μg pRSV-Rev, Addgene #14888, #12251, #12253, respectively) and transfer plasmid (2 μg) were transfected with calcium phosphate.

Medium was refreshed 12-16 h later. Virus-containing supernatant was collected 36-40 h post transfection and cleared through a 0.45-μm filter. Viral transductions were performed in the presence of 4 μg/mL polybrene (Sigma-Aldrich) at a multiplicity of infection less than 1.

### Clonogenic survival assays

DLD1 or RPE1-hTERT cells were seeded in 6-well plates (200-800 cells per well, depending on genotype). For drug sensitivity assays, treatments were applied the next day. Medium was refreshed after 6 days in all cases. At the end of the experiment, after removing medium, cells were rinsed with PBS and stained with 0.4% (w/v) crystal violet in 20% (v/v) methanol for 45 min. The stain was aspirated, and plates were rinsed twice in deionized water and air-dried. Colonies were counted using a GelCount instrument (Oxford Optronix).

### Two-color competitive growth assays

The two-color competitive growth assays were performed as previously described^67^. Briefly, cells were transduced with virus particles expressing NLS-mCherry-sgAAVS1 (control) or an NLS-GFP-sgRNA targeting a specific gene of interest (see Table S3). 24 h after transduction, virally transduced cells were selected using 20 µg/mL puromycin (Life Technologies #A1113802) for 72 h. 4d post infection, mCherry-and GFP-expressing cells were mixed 1:1 (1,000 cells each) and seeded in a 24-well plate in triplicate. Cells were imaged for GFP and mCherry signals 24 h after initial plating (t = 0) and at the indicated time points using INCell 6000 Analyzer system with a 4x objective (GE Healthcare LifeSciences). During the course of the experiment, cells were subcultured when confluency reached ∼80%. Segmentation and counting of GFP-and mCherry-positive cells were performed using a custom Acapella script (PerkinElmer)

### Immunoblotting

Cells were lysed with 2X SDS sample buffer (20% (v/v) glycerol, 2% (w/v) SDS, 0.01% (w/v) bromophenol blue, 167 mM Tris-Cl pH 6.8, 20 mM DTT) and boiled for 8 min before separating by SDS-PAGE on gradient gels (BioRad, #4561084 or #4561085). Proteins were transferred to PVDF membranes with Trans-Blot Turbo Transfer Pack (BioRad, #1704156) using a Trans-Blot Turbo Transfer System (BioRad). After blocking with 5% milk in TBST, membranes were incubated with primary antibodies at 4 °C overnight. Membranes were then washed three times for 5 min with TBST, followed by probing with appropriate secondary antibodies for 1 h at room temperature. After another three 3 x 5min washes with TBST, secondary antibody detection was achieved using an Odyssey Scanner (Li-Cor) or enhanced chemiluminescence (Thermo Fisher Scientific, SuperSignal West Pico PLUS Chemiluminescent Substrate, #34579 or West Femto Maximum Sensitivity Substrate, #34095).

### Co-Immunoprecipitation

One day before plasmid transfection by calcium phosphate, 293T cells were plated in 10-cm dishes with five million cells per dish. For TOPBP1 BRCT7/8 immunoprecipitation, 10 μg of pCIB-2xHA-DDIAS was co-transfected with 10 μg pCIB_3xFlag, pCIB-3xFlag-BRCT7/8 or pCIB-3xFlag-BRCT7/8 K1317A. For DDIAS IP, 10 μg pHIV-3xFlag, pHIV-3xFlag-DDIAS or indicated mutants were used. Cells were refreshed after 24h with culture medium containing 100 ng/mL nocodazole to enrich mitotic cells. After 16h of treatment, cells were collected and lysed by incubation on ice for 30 min in lysis buffer (50 mM Tris-HCl pH 7.5, 100 mM NaCl, 2 mM EDTA, 5 mM NaF, 0.2% NP-40, 1 mM MgCl_2_, 10% glycerol), supplemented with cOmplete EDTA-free protease inhibitor cocktail (Roche, # 11836170001) and 25 U/mL Benzonase (Sigma, # E1014). After adjusting NaCl and EDTA concentrations to 200 mM and 2 mM, respectively, lysates were then cleared by centrifugation at 16,000 x g for 15 min, followed by incubation with 15 μL of anti-FLAG M2 magnetic beads (Sigma, #M8823-5ML) for 4 hours with end-to-end mixing at 4 °C. Beads were collected using a magnetic rack and washed 4 x 5 min with 500 μL lysis buffer. Bound protein was harvested by boiling beads in 40 μL 2x SDS sample buffer and analyzed by immunoblotting.

### Split-TurboID based proximity labeling and mass spectrometry

To develop the split-enzyme proximity labeling system, we generated expression vectors that incorporated N- and C-terminal TurboID splits fused to epitope tags and exchangeable baits. FKBP and FRB domains were used as initial fusion partners to identify optimal TurboID split sites through Gateway-compatible cloning. The N-terminal construct was generated by creating an entry clone containing residues 1-97 amplified from synthesized, human codon-optimized TurboID (General Biosystems Inc., Morrisville, NC, USA) via BP cloning into pDONR223. In parallel, FKBP was PCR-amplified with primers containing KpnI/XhoI sites, a FLAG epitope tag, and a GS2 linker sequence and cloned into the Gateway-compatible pDEST pcDNA5 backbone. LR recombination generated the expression construct containing the N-terminal TurboID fragment fused to FLAG-GS2-FKBP (here after referred to as mT_N_). The C-terminal construct (mT_C_) was similarly assembled using residues 98-321 of TurboID in pDONR223 and FRB (amino acids 2031-2113 of human mTOR) with a Myc tag and GS2 linker in the pDEST backbone. Both FKBP and FRB were exchanged with genes of interest using the NheI/PacI restriction sites. All parental vector sequences were verified with confirmation digests and Sanger sequencing. The N-terminal fragment (mT_N_) of the Split-TurboID and an FKBP12-derived destabilizing domain (DD)^68^ were respectively cloned at the N-terminus and C-terminus of Flag-CIP2A to generate the plasmid pHIV-mT_N_-Flag-CIP2A-FKBP-DD. The C-terminal fragment (mT_C_) of the Split-TurboID was cloned at C-terminus of TOPBP1 in pCIB-HA2-TOPBP1. K1317A and Δ756-891 mutants of TOPBP1 were generated by site-directed mutagenesis. Using these plasmids, mT_N_-Flag-CIP2A-FKBP-DD and HA2-TOPBP1-mT_C_ derivatives were introduced into DLD1 *BRCA2*^-/-^ cells by lentiviral infection. For each condition, cells were plated in nine 15-cm dishes with 4.5 million cells per dish. The expression of mT_N_-Flag-CIP2A-FKBP-DD was induced with 1 µM AquaShield-1 the next day. After 24 h, cells were refreshed with medium containing 1 µM AquaShield-1, 100 ng/mL nocodazole and 200 µM biotin for 16h to enrich mitotic cells and enable the biotin labeling. Mitotic cells were then shaken off, pelleted and washed with ice-cold PBS. Two biological replicates were performed.

Cells were lysed in modified RIPA buffer [50 mM Tris-HCl, pH 7.5, 150 mM NaCl, 1 mM EDTA, 1 mM MgCl_2_, 1% NP40, 0.1% SDS, 0.5% sodium deoxycholate, and 1x Protease Inhibitor cocktail (Sigma-Aldrich, Cat# P8340)], sonicated and treated with TurboNuclease (BioVision, Cat#9207) and RNAse (Sigma) before incubating with streptavidin sepharose resin for 3 hours in 4°C with rotation. Subsequently, digestion with trypsin was performed on-beads. Digested peptides were analyzed using a nano-HPLC (High-performance liquid chromatography) coupled to tandem mass spectrometry (MS/MS). Briefly, peptides were reconstituted in 5% formic acid and loaded onto a 100 µm x 15cm nano-spray emitter which was generated from fused silica capillary tubing and packed with C18 reversed-phase material (Reprosil-Pur 120 C18-AQ, 3µm) resuspended in methanol. Peptides were eluted from the column with an acetonitrile gradient generated by an Eksigent ekspert nanoLC 425 and analyzed on a TripleTOF 6600 (AB SCIEX, Concord, Ontario, Canada). Tandem mass spectrometry spectra were acquired in a data-dependent mode (DDA) where the first DDA scan had an accumulation time of 250 ms within a mass range of 400-1800 Da followed by 10 MS/MS scans of the top 10 peptides identified in the first DDA scan, with accumulation time of 100 ms for each MS/MS scan.

### Protein purification

DNA coding for the BRCT7/8 domain (amino acids 1264-1493, WT or K1317A) of TOPBP1 (Uniprot ID: Q92547) with a TEV-cleavable N-terminal 6xHis-tag was cloned into pProEx-HTa (Invitrogen). *E. coli* strain BL21 DE3 RIL was transformed with the expression plasmid and 5 mL of overnight culture grown at 37°C in Lysogeny Broth was added to 1 L of Terrific Broth containing 100 mg/L ampicillin. The culture was grown at 37°C with shaking at 180 rpm until an optical density of 1.2 was reached. The temperature was lowered to 16°C and IPTG was added to a final concentration of 0.5 mM. The culture was incubated overnight and the bacterial pellet was harvested by centrifugation at 6000g. The harvested bacterial pellet was then lysed by homogenization (Avestin) in buffer containing 50 mM HEPES pH 7.5, 500 mM NaCl, 1 mM TCEP, 5 mM imidazole. The lysate was clarified by centrifugation and passed through 1 ml HisTrap HP column (Cytiva). The column was washed with 50 ml of lysis buffer and bound protein was eluted using a gradient of imidazole. Pooled fractions were concentrated and then injected onto a Superdex 75 16/600 column (Cytiva) equilibrated with a buffer containing 20 mM HEPES pH 7.5, 400 mM NaCl and 1 mM TCEP for final polishing and buffer exchange. Pooled fractions were concentrated to 14.87 mg/mL and flash frozen in liquid nitrogen.

Wild type and mutant DDIAS OB domain (residues 2-197) peptides were expressed as N-terminal maltose-binding protein (MBP) tagged fusion peptides in *E. coli* BL21 (DE3) cells cultured in Terrific Broth. Expression was induced by adding IPTG to a final concentration of 0.3 M to growth medium for ∼18 h at 18 °C. Cells were harvested by centrifugation, resuspended in lysis buffer [50 mM Tris (pH 7.5), 500 mM NaCl, 1 mM PMSF, 1 mM DTT], homogenized and sonicated. Cleared lysates were incubated with 10 ml of amylose resin (New England Biolabs, #E8021L) to capture the fusion peptide, which was then eluted with 50 mM maltose. Fractions containing the MBP-tagged DDIAS-OB peptide were pooled, concentrated and loaded onto a Superdex S200 (Cytiva) sizing column equilibrated in the lysis buffer. A total of fifty 500 µl fractions were collected; fractions containing the desired peptide peaks were pooled together, concentrated, snap frozen in liquid nitrogen and stored at -80 °C.

### Fluorescence polarization assay

N-terminally FITC-labeled peptides (Biomatik) listed below were dissolved in DMSO and diluted in binding buffer (20 mM HEPES pH 7.5, 150 mM NaCl, 1 mM TCEP, 0.01% Brij-35) to a concentration of 25 nM. Purified TOPBP1 BRCT7/8 WT and K1371A were diluted in binding buffer to generate a 450 µM stock and serially diluted to the desired concentration. 25 µL each of purified protein and peptide were mixed in a 384-well microplate (Corning, #3573), and fluorescent polarization was measured with BioTek Synergy microplate reader.

pBRIP1: FITC-ESIYFpTPELYDPEDT-NH_2_ DDIAS S676: FITC-SYDASADLFDDIAK-NH_2_ DDIAS pS676: FITC-SYDApSADLFDDIAK-NH_2_ DDIAS S993: FITC-ETRSAWSPELFS-NH_2_ DDIAS pS993: FITC-ETRSAWpSPELFS-NH_2_

### Electrophoretic Mobility Shift Assay (EMSA)

DNA binding of recombinant DDIAS-OB domain was tested with EMSA using 5’-FAM-labelled ssDNA and dsDNA (completely or partially hybridized) probes. Probe hybridization was achieved by mixing ssDNA probes in hybridization buffer [50 mM Tris (pH 7.5), 250 mM NaCl] which was then heated at 80° C for 10 min, followed by gradual cooling overnight. For electrophoretic mobility shift assays, 10 nM DNA probe was incubated with increasing concentrations of purified recombinant peptide, in the presence of 1% 1mg/ml BSA in a final concentration of 150 µM/µl NaCl (10 µl reaction volume). Reactions were carried out in the dark at room temperature for ∼30 min. Subsequently, 2 µl of 50% glycerol was added to the reaction mixture and resolved on a 6% native polyacrylamide-TBE gel. The gel was scanned on a Typhoon FLA 9500 biomolecular imager (GE Healthcare), and band intensity was processed using Image Studio lite v5.2 (Li-Cor Biosciences, Lincoln, NE).

### Time-resolved fluorescence energy transfer (TR-FRET)

For TR-FRET assays, increasing concentration (0-1 µM) of purified MBP-tagged recombinant DDIAS-OB fold (wild type and mutant) was incubated with 5’-FAM-labelled ssDNA (10 nM) in the presence of a Lumi4 Tb-conjugated anti-MBP mouse IgG1 antibody (Revvity; 61MBPTAA) in low-volume 96 well plates (Revvity, 66PL96005). The final reaction mixture contained 1 mg/ml BSA and 150 µM/µl NaCl in PPI-Terbium detection buffer (Revvity, 61MBPTAA). For competition assays, the concentration of purified MBP-DDIAS OB-fold protein and labelled ssDNA was kept constant at 0.5 µM and 10 nM, respectively, whereas the cold probe was titrated from 0-10 µM. The reaction was allowed to proceed for ∼30 min at room temperature in the dark. With a BioTek Synergy Neo plate reader, the reactions was excited at 340 nm, and the emission was recorded at 495 nm and 520 nm, with a 60 µsec delay in between. OD_525_ values were divided by OD_495_ values and normalized to the no protein control.

### Immunofluorescence

Cells were grown and fixed on glass coverslips with 3% PFA, permeabilized with 0.3% Triton X-100 in PBS, and blocked with 5% BSA in PBS with 0.2% Tween-20. For RPA staining, cells were pre-extracted with 0.5% Triton X-100 in CSK buffer (10 mM 1,4-piperazinediethanesulfonic acid (PIPES), pH 7.0, 100 mM NaCl, 300 mM sucrose, 3 mM MgCl_2_, 1mM EGTA) for 15 min on ice, washed once with CSK buffer and once with PBS before PFA fixation. Cells were then stained for 2 h with primary antibodies in blocking buffer, washed three times with PBS + 0.2% Tween-20, incubated for 1 h with appropriate secondary antibodies in blocking buffer plus 0.8 ug/ml DAPI, then washed twice with PBS + 0.2% Tween-20 and a final wash with PBS. Coverslips were mounted onto glass slides with ProLong Gold mounting reagent (ThermoFisher Scientific, # P36930). Images were acquired using a Zeiss LSM780 laser-scanning microscope (Zeiss) with a 63×, 1.4-numerical aperture Plan-Apochromat oil-immersion objective or Nikon BioPipeline (Nikon Instruments Inc.) with a 60×, 1.2-numerical aperture Plan-Apochromat water-immersion objective. Foci were quantified by Fiji (v2.16.0/1.54p).

### DNA fluorescent *in situ* hybridization (FISH)

DNA FISH was performed as previously described^69^. Following IF staining, cells were fixed with methanol/acetic acid (3:1) for 15 min at room temperature. Slides were rinsed with 80% ethanol and left to air dry for 5 min. Human Y-chromosome FISH probes (MetaSystems, # D-0324-100-FI) were applied to cells on coverslips, followed by denaturation at 75°C for 3 min and incubation at 37°C overnight in a humidified chamber. Slides were then submerged in prewarmed 75°C 0.4x SSC solution for 2 mins and rinsed in 2x SSC/0.05% Tween-20. After washing with PBS, cells were stained with DAPI for 10 min and rinsed with water. Coverslips were mounted onto new glass slides with ProLong Gold mounting reagent. IF-FISH images were acquired using a DeltaVision Elite microscope system. Deconvolved maximum intensity projections were generated using softWoRx (v6.0).

### Micronuclei (MN) detection

For detection of MN, cells were fixed with 3% PFA, washed 3× with PBS, permeabilized with 0.3% Triton X-100 in PBS for 10 min, washed 3× with PBS, and incubated for 1 h with PBS + DAPI (0.5 μg/mL). After the last wash with PBS, images were acquired on an INCell Analyzer 6000 automated microscope (GE Life Sciences) with a 60x objective. For each well in a 96-well plate, 100 images were acquired. MN were automatically detected and counted using the Columbus analysis tool (PerkinElmer).

### CRISPR screens

CRISPR screens were carried out as previously described^70^. Briefly, DLD1 wild type (WT) and *DDIAS*^-/-^ cells were transduced with the lentiviral TKOv3 library at a low MOI (∼0.25). Puromycin-containing medium was added the next day for selection. Three days after infection, which was considered the initial time point (t0), cells were pooled together and divided into two sets of technical replicates. Cells were subcultured every 3 d for a total of 18 d before harvesting. Screens were performed with a theoretical library coverage of ≥400 cells per sgRNA maintained at every step. Genomic DNA was isolated using the QIAamp Blood Maxi Kit (Qiagen) and genome-integrated sgRNA sequences were amplified by PCR using NEBNext Ultra II Q5 Master Mix (New England Biolabs). i5 and i7 multiplexing barcodes were added in a second round of PCR and final gel-purified products were sequenced on an Illumina NextSeq500 system at the LTRI NBCC facility (https://nbcc.lunenfeld.ca/) to determine sgRNA representation in each sample. The screen results were analyzed using BAGEL2^47^.

## Data deposition

The CRISPR screen sequencing was deposited at the NCBI with the Project ID: PRJNA1272578. Proteomics data has been deposited as a complete submission to the MassIVE repository (https://massive.ucsd.edu/ProteoSAFe/static/massive.jsp) and assigned the accession number MSV000098110. The dataset is currently available for reviewers at ftp://MSV000098110@massive.ucsd.edu. Please login with username MSV000098110_reviewer; password: cg5cRWWHEg2LNAiQ. Finally, all Source Data is available for reviewers at Zenodo at this <u>link</u>.

---

**Figure S1.**
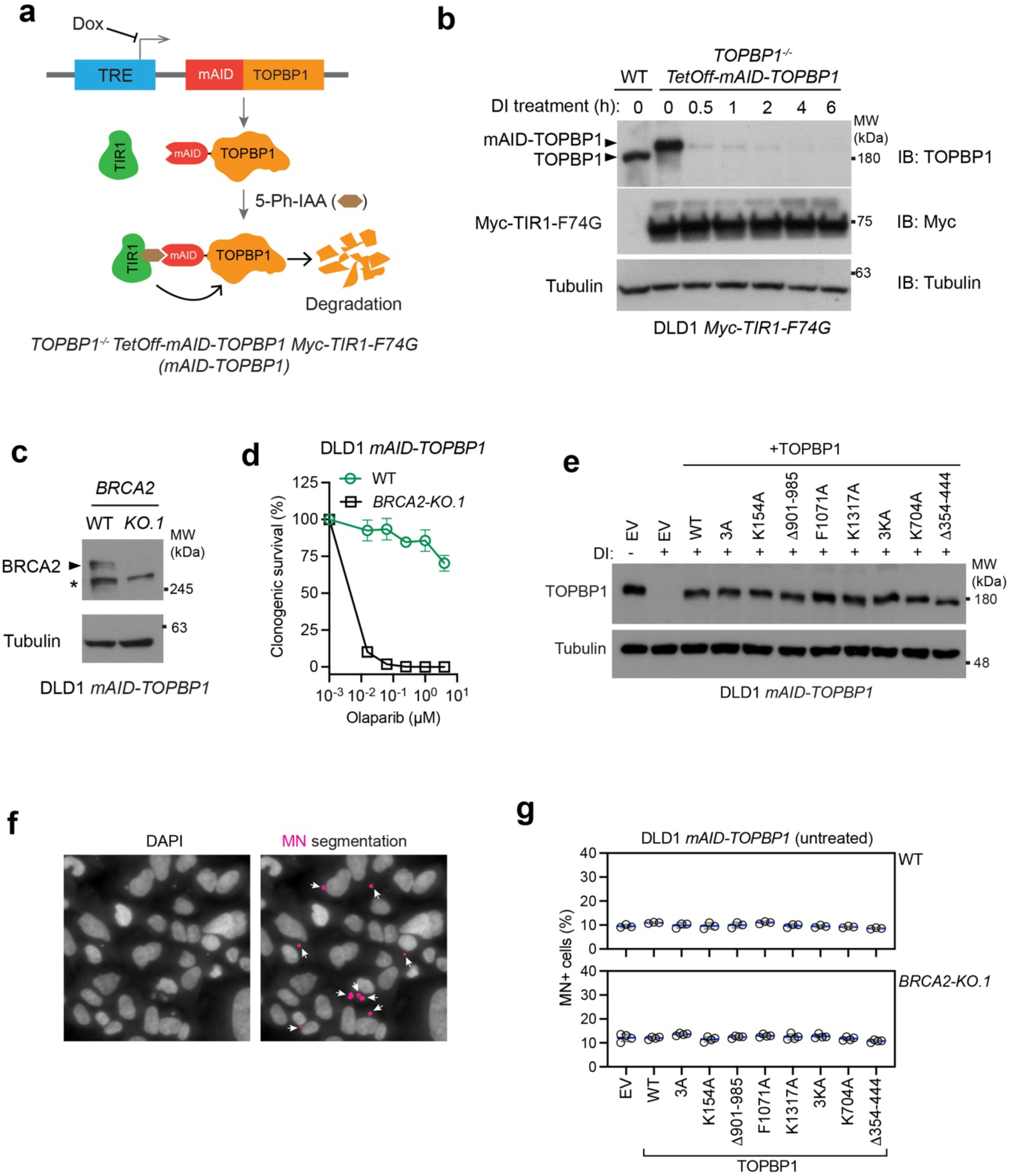
A conditional gene replacement system for TOPBP1 in DLD1 cells. Relates to. Figure 1**.a**) Schematic showing the conditional silencing strategy for TOPBP1. (**b**) Immunoblot showing the knockout (KO) of endogenous *TOPBP1* and dynamics of mAID-TOPBP1 depletion by DI (doxycycline (1 μg/mL) + 5-Ph-IAA (1 μM)). (**c**) Immunoblot verifying BRCA2 KO in DLD1 *mAID-TOPBP1* cells. (**d**) Clonogenic survival of DLD1 *mAID-TOPBP1* WT and *BRCA2*^-/-^ cells in response to olaparib. Bars indicate the mean ± SD. n=3 independent experiments. (**e**) Immunoblot showing the expression of *TOPBP1* variants in DLD1 *mAID-TOPBP1* cells treated with DI for 8h. (**f**) Representative images of micronuclei segmentation in DLD1 *mAID-TOPBP1* cells by Columbus software. (**g**) Quantification of micronuclei in DLD1 *mAID-TOPBP1* WT or *BRCA2*^-/-^ cells expressing the indicated *TOPBP1* variants or EV (empty vector) without DI treatment. Bars indicate the mean ± SD. n=3 independent experiments for WT cells and n= 4 for *BRCA2*^-/-^ cells. Tubulin, loading control.

**Figure S2.**
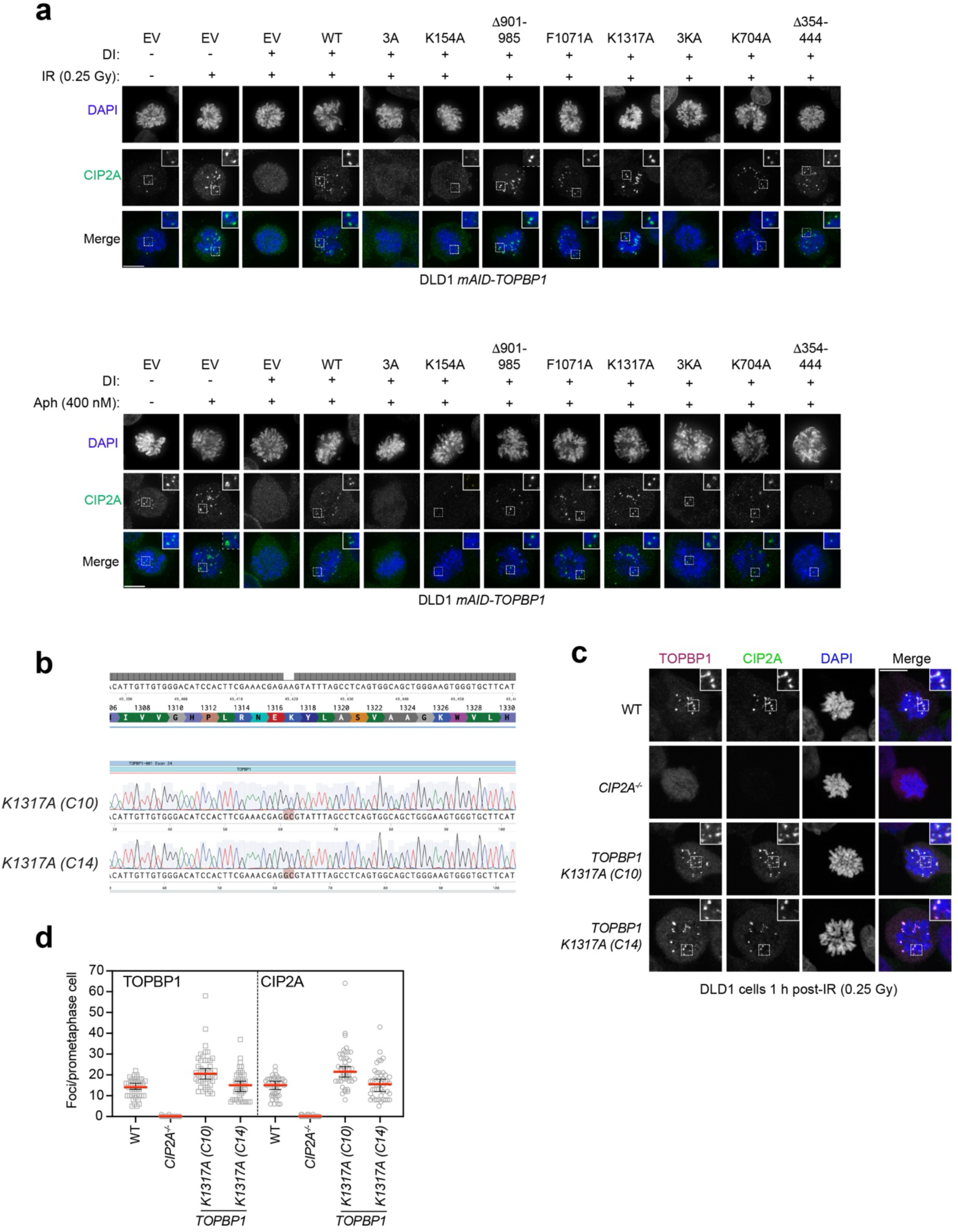
Impact of TOPBP1 mutants on the localization of CIP2A to mitotic DNA lesions. Relates to. Figure 1. (**a**) Representative micrographs of the experiment shown in Fig. 1e. Treatments: upper panels: 1h after 0.25 Gy IR; lower panels: 16h of 400nM aphidicolin (Aph) (**b**) Sequencing chromatograms verifying the *TOPBP1 K1317A* mutation in two DLD1 clones. (**c**) Quantification of CIP2A and TOPBP1 foci in mitotic DLD1 cells with the indicated genetic background after IR exposure. Blue dots represent measurements from individual cells. Bars indicate the mean ± SD. n=3 independent experiments. (**d**) Representative micrographs of the experiment shown in **c**. Scale bar, 10 μm.

**Figure S3.**
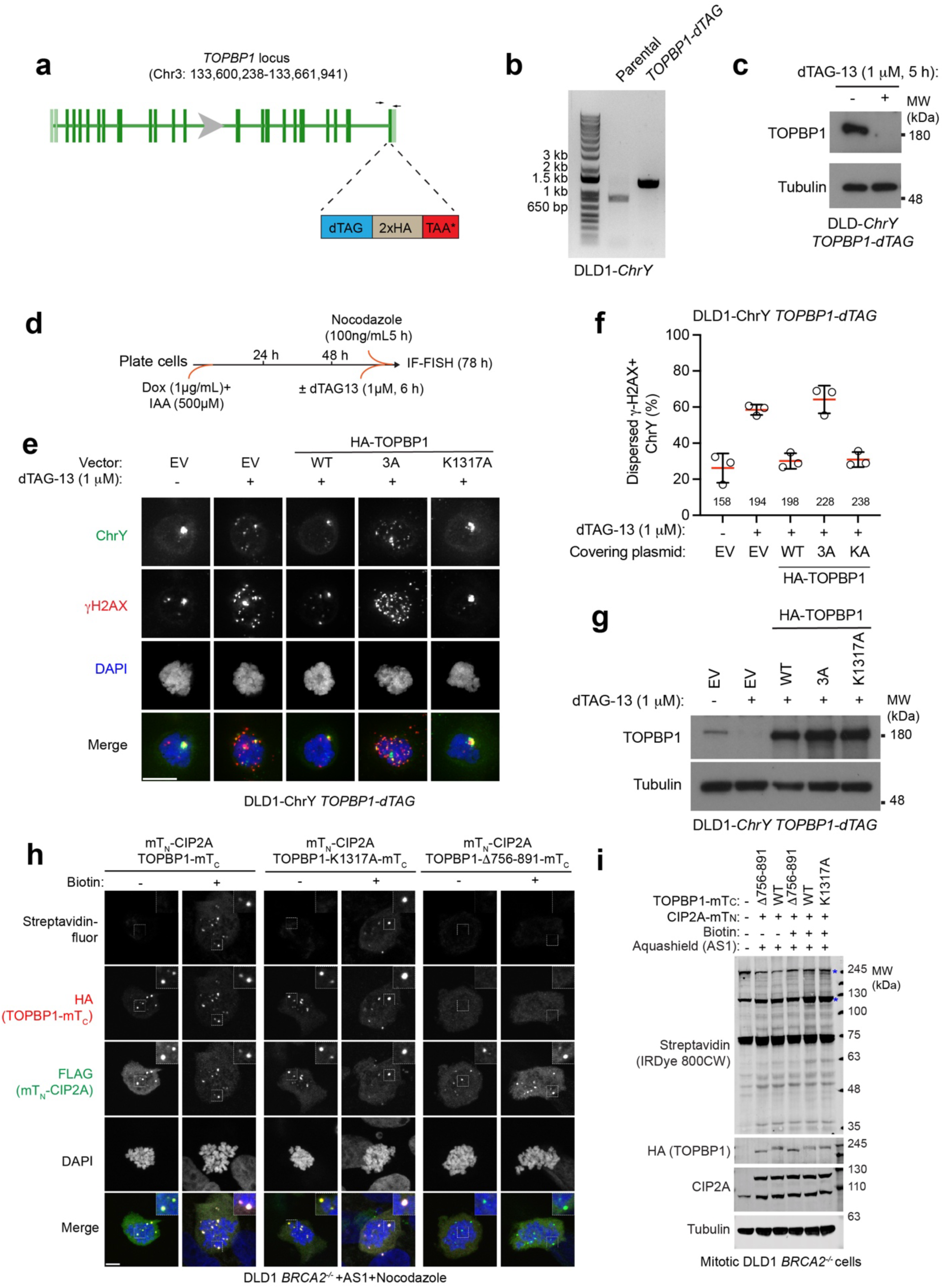
The identification of DDIAS as a CIP2A-TOPBP1 complex-binding protein. Related to. Figure 2. (**a**) Schematic showing endogenous tagging with dTAG-2xHA at the C-terminus of TOPBP1 in DLD1 ChY cells. (**b**) Agarose gel image of PCR products from indicated cells verifying successful bi-allelic tagging with the primers indicated in **a**. (**c**) Immunoblot showing degradation of TOPBP1-dTAG-2xHA by dTAG-13 in DLD1-ChY *TOPBP1-dTAG* cells. (**d**) Experimental outline for **e, f**. (**e**) Representative micrographs showing the status of the micronucleated Y-chromosome in mitotic cells from the experiment in **f**. (**f**) Quantitation of dispersed Y-chromosome fragments in mitotic DLD1-ChY *TOPBP1-dTAG* cells expressing the indicated *TOPBP1* variants or EV after depletion of endogenous TOPBP1. Numbers of cells analyzed are for each condition shown. Bars indicate the mean ± SD. n=3 independent experiments. KA: K1317A. (**g**) Immunoblot confirming expression of the indicated *TOPBP1* variants in DLD1-ChY *TOPBP1-dTAG* cells. (**h**) Immunofluorescence analysis verifying the biotinylation activity of Split-TurboID in DLD1 BRCA2^-/-^ cells expressing CIP2A-FKBP^DD^ and the indicated variants of TOPBP1 respectively tagged with the N-terminus and C-terminus of miniTurbo (mT_N_ and mT_C_). FKBP^DD^ is an FKBP12-derived destabilization domain (DD) that was fused to C-terminus of mT_N_-CIP2A to enable its inducible expression (see Methods for detail). mT_N_-CIP2A-FKBP^DD^ expression was induced by 1µM AquaShield-1 (AS1) for 24h. Mitotic cells were enriched by nocodazole (100ng/mL) in the presence or absence of 200 μM Biotin for 16h. Note that the presence of CIP2A foci in the cells that express the TOPBP1 Δ756-891 mutant is explained by the fact that endogenous TOPBP1 is present. (**i**) Immunoblot verifying the biotinylation activity of Split-TurboID in DLD1 BRCA2^-/-^ cells expressing mT_N_-CIP2A-FKBP^DD^ and indicated variants of TOPBP1-mT_C_. Asterisks indicate self-biotinylation. Scale bar, 5 μm. Tubulin, loading control.

**Figure S4.**
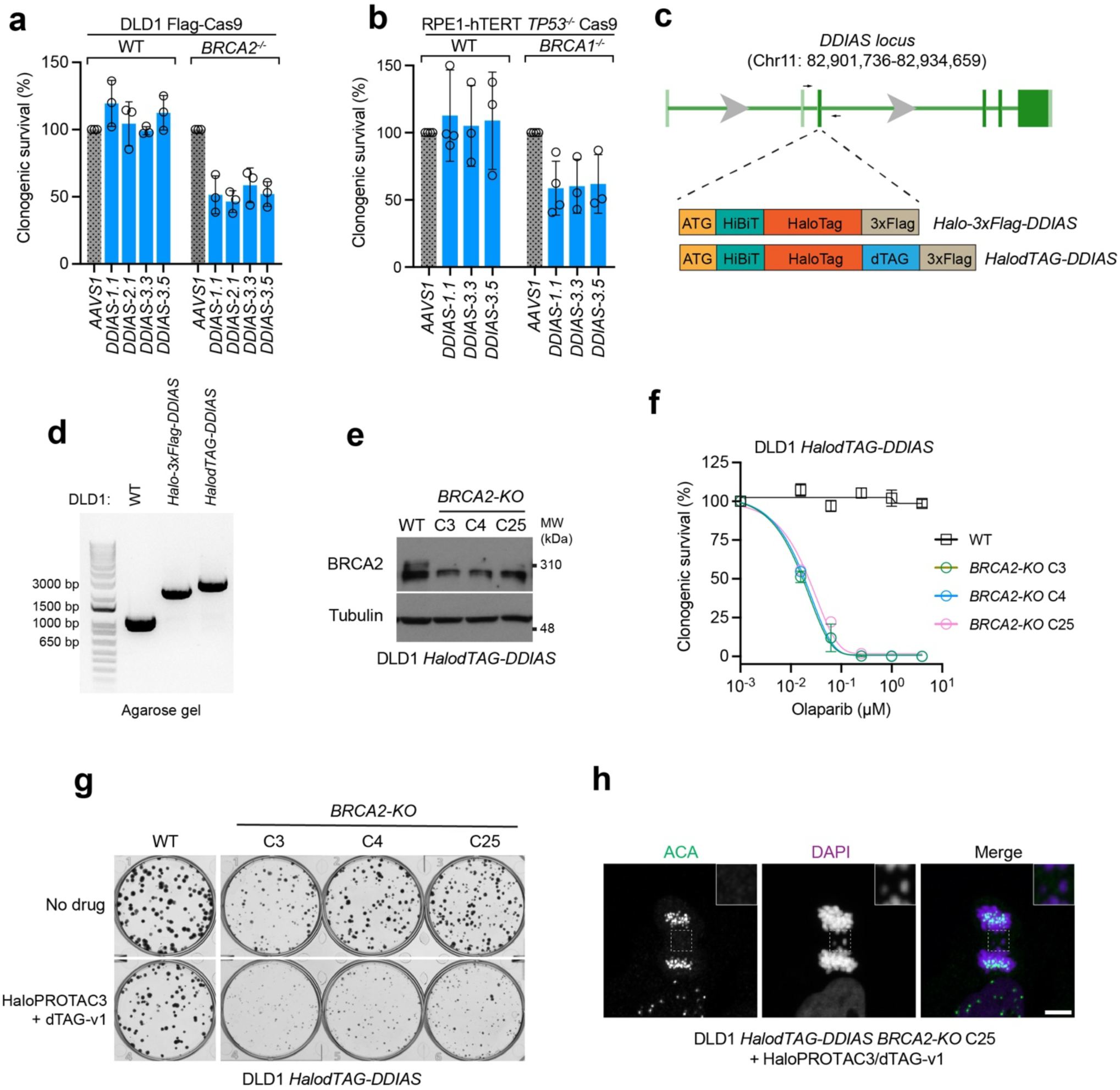
DDIAS promotes the viability of HR-deficient cells. Relates to. Figure 2. (**a, b**) Clonogenic survival of DLD1 Cas9 (WT or *BRCA2*^-/-^) (**a**) and RPE1-hTERT TP53^-/-^ Cas9 (WT or *BRCA1*^-/-^) (**b**) cells expressing the indicated *DDIAS* or *AAVS1*-targeting sgRNAs. Bars indicate the mean ± SD. n ≥3 independent experiments. (**c**) Schematic showing endogenous tagging of HiBit-HaloTag-3xFlag or HiBit-HaloTag-dTAG-3xFlag at the N-terminus of DDIAS. (**d**) Agarose gel image of PCR products from indicated cells verifying successful bi-allelic tagging with the primers indicated in (**c**). (**e**) Immunoblot verifying the knockout of *BRCA2* in DLD1 *HalodTAG-DDIAS* cells. Tubulin, loading control. (**f**) Clonogenic survival of DLD1 *HalodTAG-DDIAS* WT and *BRCA2*^-/-^ cells in response to olaparib. Bars indicate the mean ± SD. n=3 independent experiments. (**g**) Representative images of colony forming from of the experiment shown in Fig 2f. (**h**) Representative micrographs showing acentric lagging chromosomes in DLD1 *HalodTAG-DDIAS BRCA2*^-/-^ cells with DDIAS depletion as shown in Fig 2i. Scale bar, 5 μm.

**Figure S5.**
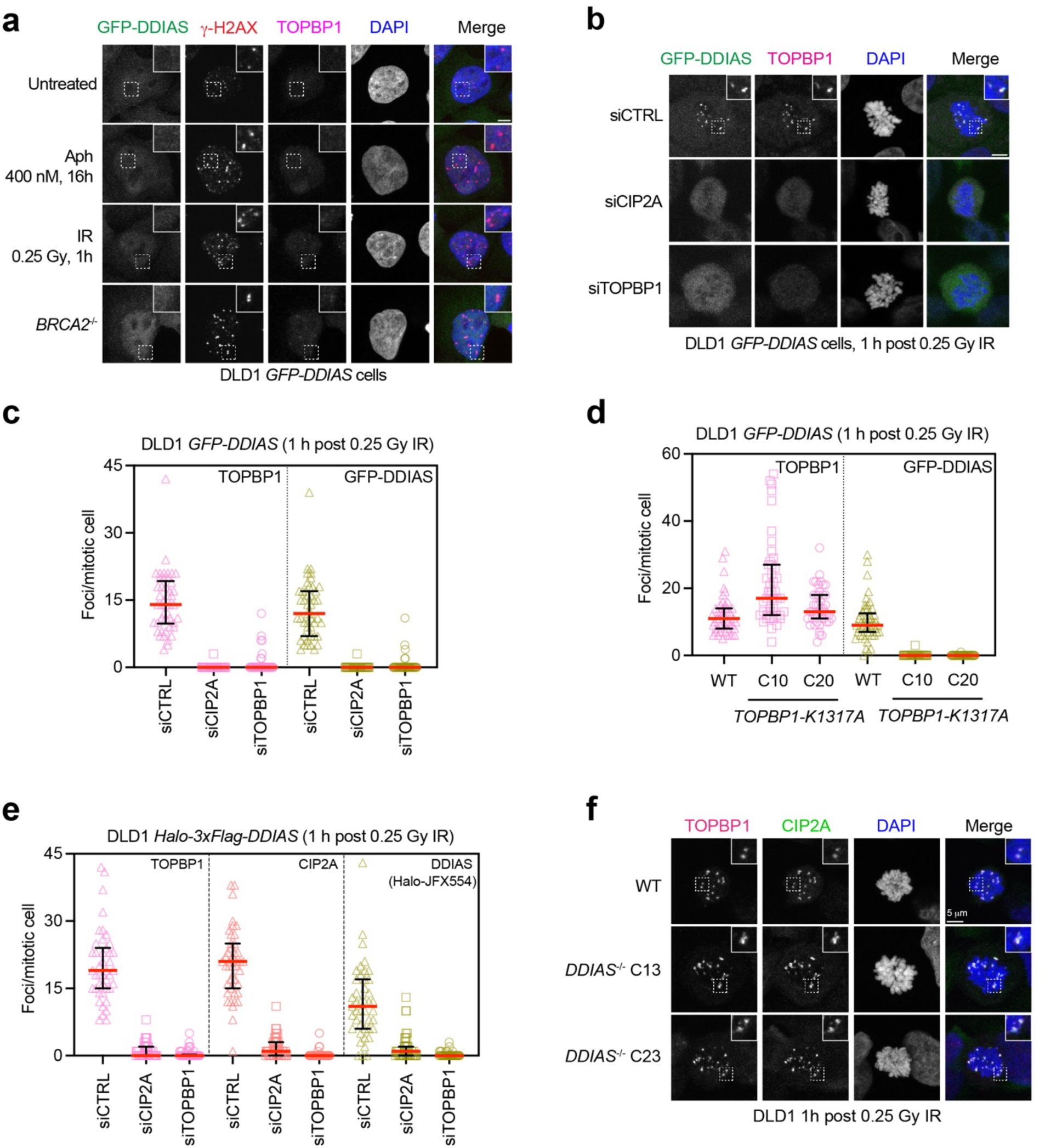
Additional data on the localization of DDIAS at mitotic DNA lesions. Relates to. Figure 3. (**a**) Representative micrographs of Fig. 3b showing the localization of GFP-DDIAS in interphase DLD1 cells under different DNA damaging conditions. (**b**) Immunofluorescence analysis of TOPBP1 and GFP-DDIAS foci in mitotic DLD1 cells after siRNA-mediated depletion of CIP2A or TOPBP1 followed by IR treatment. (**c**) Quantification of TOPBP1 and GFP-DDIAS foci in individual mitotic cells from the experiment shown in (**b**). Bars indicate the mean ± SD. n=3 independent experiments. (**d**) Quantification of TOPBP1 and GFP-DDIAS foci in individual mitotic cells from the experiment shown in Fig. 3c. Bars indicate the mean ± SD. n=3 independent experiments. (**e**) Quantification of TOPBP1, CIP2A and endogenous Halo-DDIAS foci in individual mitotic cells from the experiment shown in Fig. 3d. Bars indicate the mean ± SD. n=3 independent experiments. (**f**) Representative micrographs from the experiment in Fig. 3e showing CIP2A and TOPBP1 foci in mitotic DLD1 WT or *DDIAS*^-/-^ cells after IR treatment. Scale bar, 5 μm.

**Figure S6.**
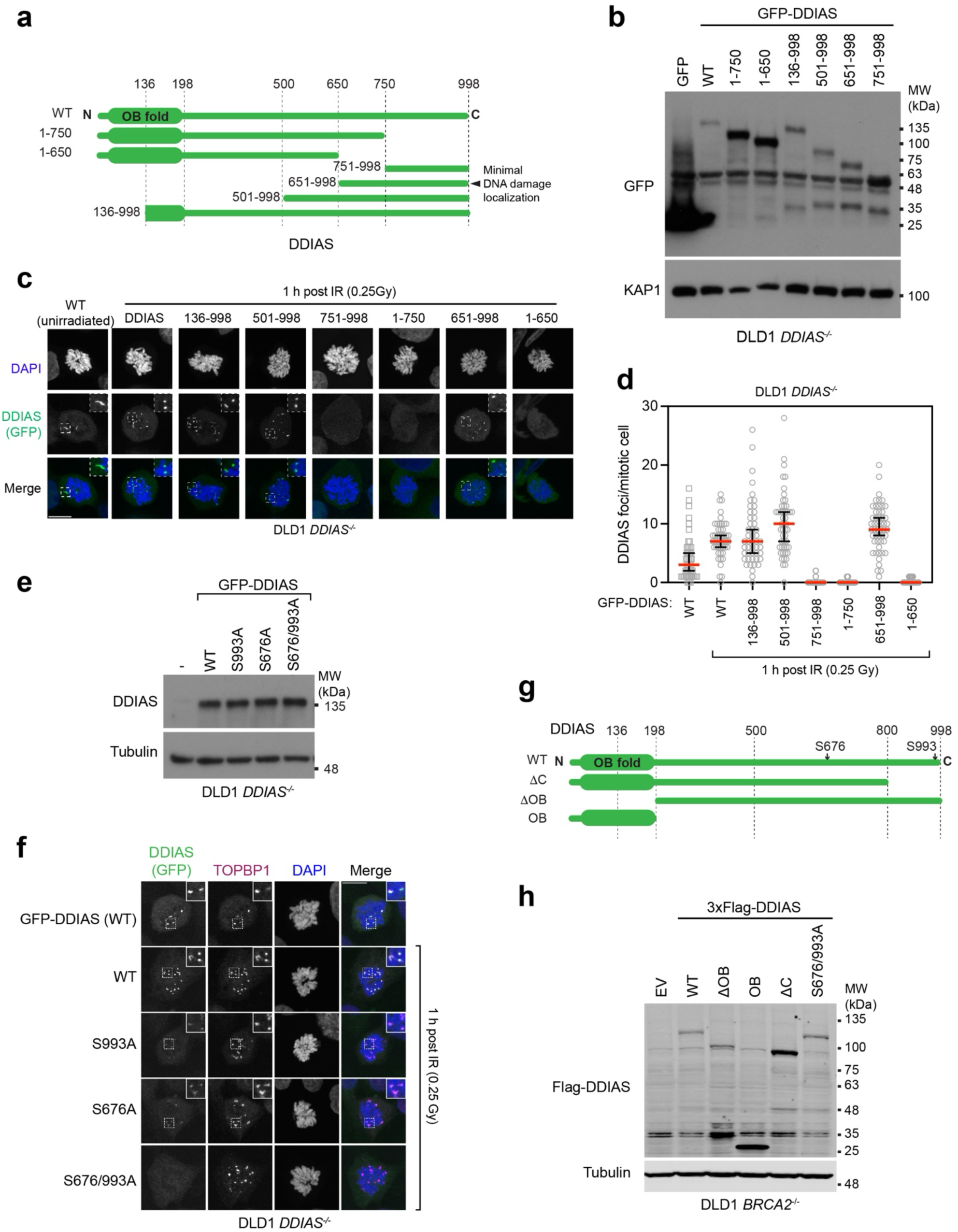
DDIAS residues S676 and S993 mediate its recruitment to DNA damage sites. Relates to. Figure 4. (**a**) Schematic showing the domain structure of WT and the indicated variants of DDIAS. (**b**) Immunoblot showing the expression of WT and the indicated variants of GFP-DDIAS in DLD1 *DDIAS*^-/-^ cells. KAP1, loading control. (**c**) Immunofluorescence analysis showing the localization of WT and indicated variants of DDIAS in mitotic DLD1 *DDIAS*^-/-^ cells after IR treatment. (**d**) Quantification of WT and mutated GFP-DDIAS foci in individual mitotic cells from the experiment shown in (**c**). Bars indicate the mean ± SD. n=3 independent experiments. (**e**) Immunoblot showing the expression of WT and the indicated variants of GFP-DDIAS in DLD1 *DDIAS*^-/-^ cells. Tubulin, loading control. (**f**) Representative micrographs of the experiment presented in Fig. 4c showing TOPBP1 and GFP-DDIAS foci in mitotic DLD1 *DDIAS*^-/-^ cells after IR treatment. (**g**) Schematic showing the domain structure of WT and the indicated variants of DDIAS. (**h**) Immunoblot showing the expression of WT and the indicated sgRNA-resistant variants of 3xFlag-DDIAS in DLD1 BRCA2^-/-^ cells. Tubulin, loading control. Scale bar, 10 μm.

**Figure S7.**
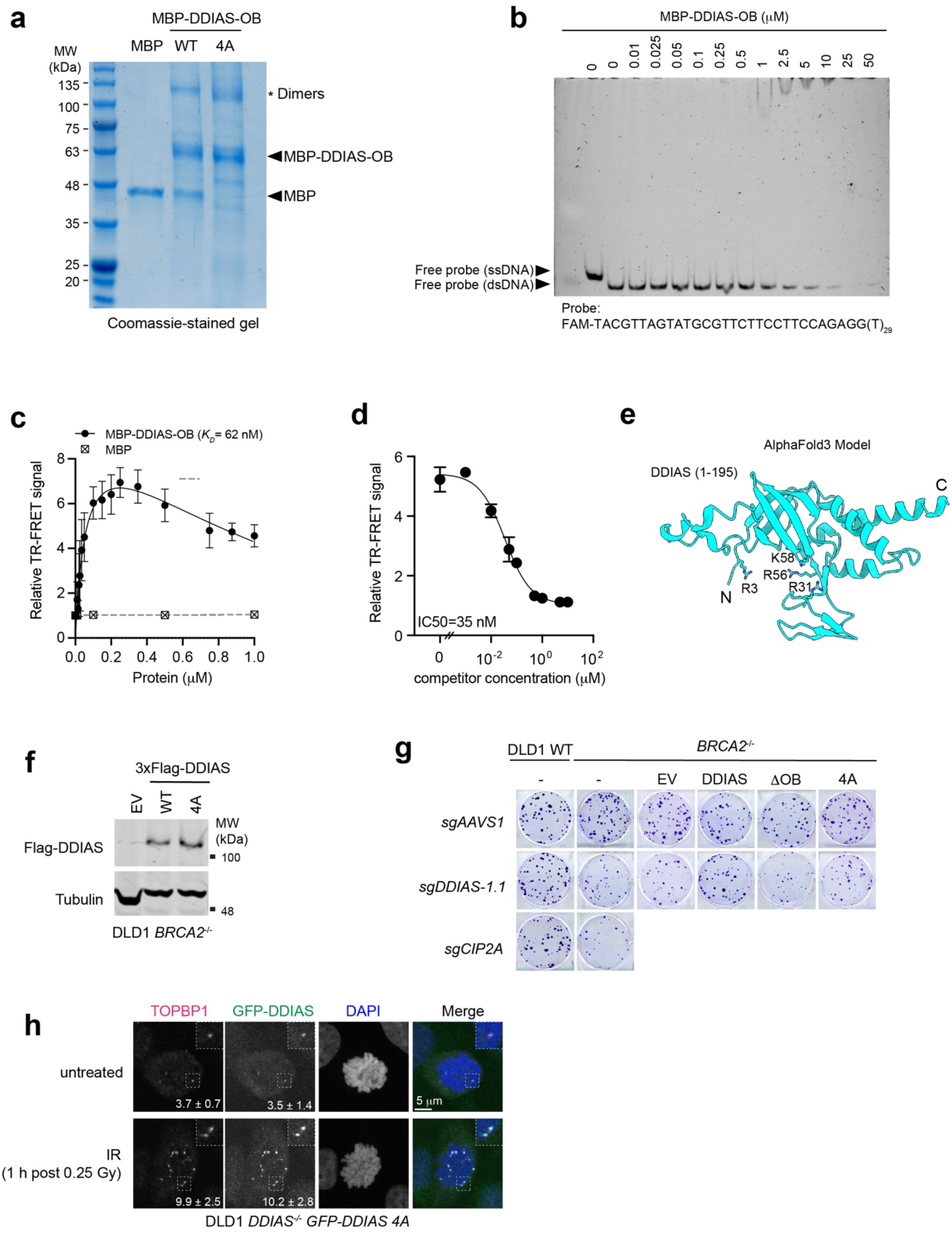
The OB-fold of DDIAS binds to ssDNA. Relates to. Figure 4. (**a**) Coomassie-stained gel of recombinant MBP, WT and 4A mutated MBP-DDIAS-OB. Purified MBP-DDIAS-OB appears to contain both monomer and dimer forms. (**b**) EMSA using recombinant MBP-DDIAS-OB and fluorescently labeled double-stranded DNA (dsDNA). n=3 independent experiments done with at least two different protein preps. (**c**) TR-FRET assay using fluorescently labeled ssDNA and increasing concentration of recombinant MBP-DDIAS-OB. Bars indicate the mean ± SD. n=3 independent experiments. (**d**) TR-FRET assay using recombinant MBP-DDIAS-OB and fluorescently labeled ssDNA with increasing concentration of unlabeled competitor. Bars indicate the mean ± SD. n=3 independent experiments. (**e**) AlphaFold3 prediction of DDIAS OB-fold (1-195) with 4 positively charged residues highlighted. (**f**) Immunoblot showing the expression of WT and 4A mutated DDIAS in DLD1 *BRCA2*^-/-^ cells. Tubulin, loading control. (**g**) Representative images of clonogenic survival experiment shown in Fig 4i. (**h**) Immunofluorescence analysis of GFP-DDIAS-4A localization in mitotic DLD1 *DDIAS*^-/-^ cells with or without IR treatment. Average numbers of GFP-DDIAS-4A foci from 3 independent experiments are indicated. Scale bar, 5 μm.

**Figure S8.**
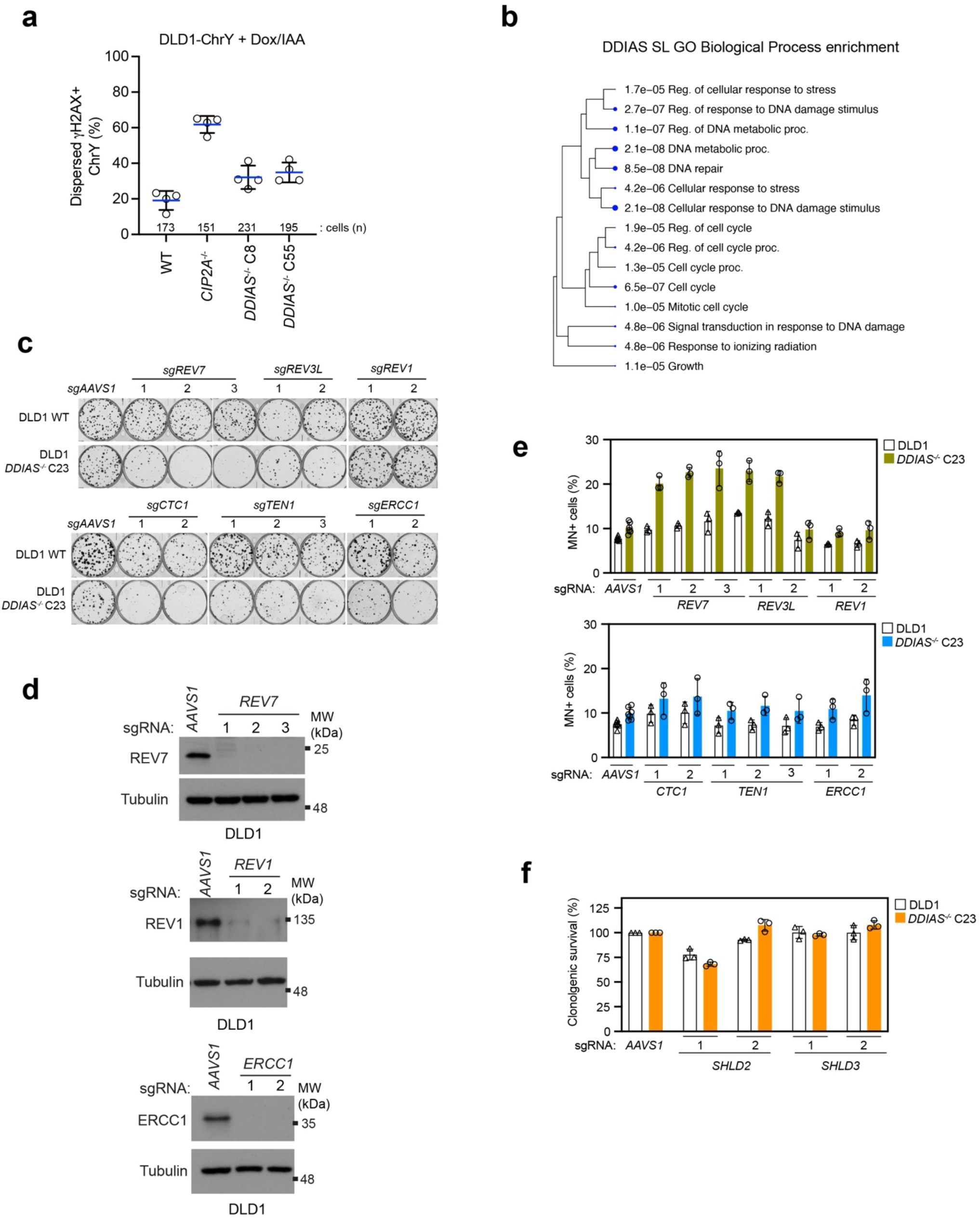
DDIAS loss is synthetic lethal with genetic loss of Pol(- and CST-coding genes. Related to. Fig 5. (**a**) Quantitation of dispersed Y-chromosome fragments in mitotic DLD1-ChY cells with the indicated genetic background. Y-chromosome micronuclei were induced as in Fig S3f. Bars indicate the mean ± SD. n=3 independent experiments. (**b**) Gene ontology enrichment analysis for hits from the DDIAS synthetic lethal screen. (**c**) Representative images of clonogenic survival experiment showed in Fig 5b. (**d**) Immunoblot verifying the efficiency of the indicated sgRNAs used in the experiment shown in Fig 5b. (**e**) Quantification of micronuclei in DLD1 WT and *DDIAS*^-/-^ cells expressing the indicated sgRNAs. Bars indicate the mean ± SD. n=3 independent experiments. (**f**) Clonogenic survival of DLD1 WT and *DDIAS*^-/-^ cells expressing sgRNAs against SHLD2 or SHLD3. Bars indicate the mean ± SD. n=3 independent experiments.

**Figure S9.**
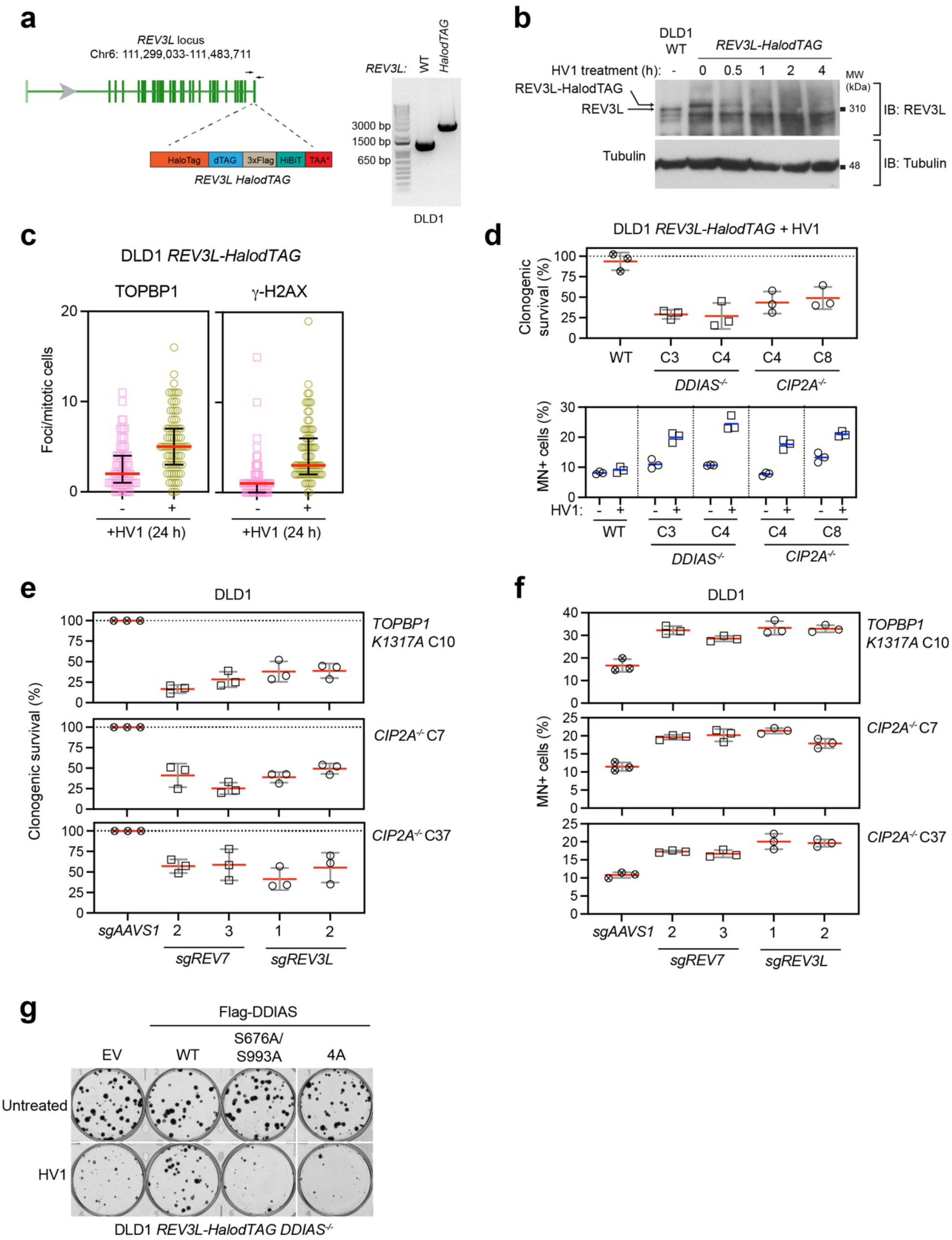
Pol(-deficient cells require for the CIP2A-TOPBP1-DDIAS pathway for genome stability. Related to. Fig 5. (**a**) Endogenous tagging of REV3L with HaloTag-dTAG-3xflag-HiBiT (HalodTAG) at the C-terminus in DLD1 cells. Agarose gel image of PCR products from indicated cells verifying successful bi-allelic tagging with the primers indicated. (**b**) Immunoblot showing REV3L degradation dynamics in DLD1 *REV3L*-*HalodTAG* cells by HaloPROTAC3 (1 μM) and dTAG-v1 (1 μM). Tubulin, loading control. (**c**) Immunofluorescence analysis of TOPBP1 and γ-H2AX foci in mitotic DLD1 *REV3L*-*HalodTAG* cells with or without REV3L depletion for 24h. Bars indicate the mean ± SD. n=4 independent experiments. (**d**) Quantification of clonogenic survival and micronuclei in DLD1 *REV3L-HalodTAG* cells and the indicated derivatives with or without REV3L depletion. Clonogenic survival data was normalized to untreated condition. Bars indicate the mean ± SD. n=3 independent experiments. (**e, f**) Quantification of clonogenic survival (**e**) and micronuclei (**f**) in DLD1 *CIP2A*^-/-^ and *TOPBP1 K1317A-*mutated cells expressing sgRNAs against *REV7* or *REV3L*. Bars indicate the mean ± SD. n=3 independent experiments. (**g**) Representative images of the clonogenic survival experiment showed in Fig 5c.

**Figure S10.**
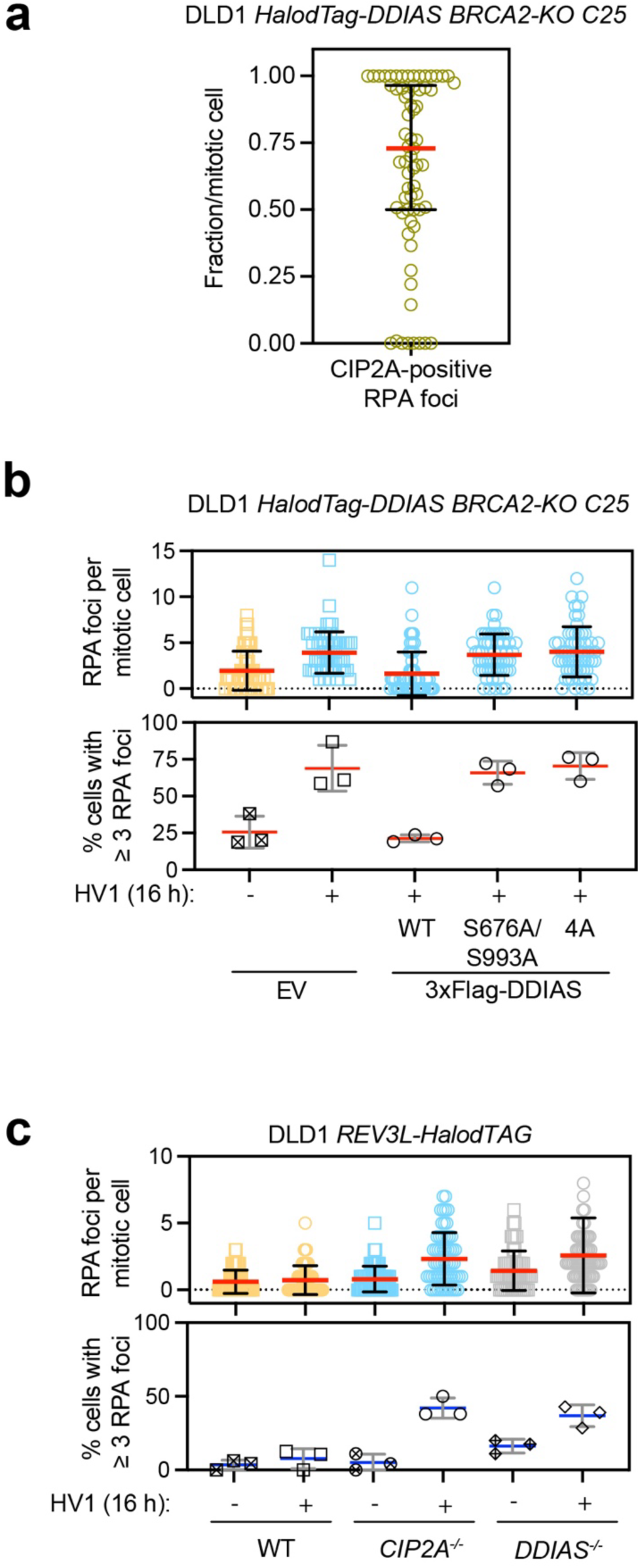
DDIAS suppresses ssDNA in mitosis. Related to. Fig 5. (**a**) Colocalization analysis of RPA and CIP2A foci in mitotic DLD1 *HalodTAG-DDIAS BRCA2*^-/-^ cells with DDIAS depletion. Bars indicate the mean ± SD. n=3 independent experiments. (**b**) Immunofluorescence analysis of RPA foci in mitotic DLD1 *HalodTAG-DDIAS BRCA2*^-/-^ cells expressing the indicated variants of DDIAS with or without depletion of endogenous DDIAS. Bars indicate the mean ± SD. n=3 independent experiments. (**c**) Immunofluorescence analysis of RPA foci in mitotic DLD1 *REV3L-HalodTAG* cells and indicated derivatives with or without REV3L depletion. Bars indicate the mean ± SD. n=3 independent experiments.

**Supplementary Table 1.** Information of cell lines and their genetically engineered derivatives used in this study.

**Supplementary Table 2.** Mass spectrometry results of CIP2A-TOPBP1 Split-TurboID experiment in DLD1 *BRCA2^-/-^* cells.

**Supplementary Table 3.** Information of sgRNA used in this study.

**Supplementary Table 4.** Results of DDIAS synthetic lethal screen in DLD1 cells.

